# Transcriptional dissection of symptomatic profiles across the brain of men and women with depression

**DOI:** 10.1101/2023.04.21.537733

**Authors:** Samaneh Mansouri, André M Pessoni, Arturo M Rivera, Carol A. Tamminga, Eric Parise, Gustavo Turecki, Eric J. Nestler, Ting-Huei Chen, Benoit Labonté

**Author notes:** Corresponding Authors: Benoit Labonté Associate professor, Department of Psychiatry and Neurosciences, Faculty of Medicine, Université Laval, Québec City, QC, Canada.

## Abstract

Major depressive disorder (MDD) is one of the most important causes of disability worldwide. While recent work provides insights into the molecular alterations in the brain of patients with MDD, whether these molecular signatures can be associated with the expression of specific symptom domains in males and females remains unclear. Here, we identified sex-specific gene modules associated with the expression of MDD, combining differential gene expression and co-expression network analyses in six cortical and subcortical brain regions. Our results show varying levels of network homology between males and females across brain regions, although the association between these structures and the expression of MDD remains highly sex-specific. We refined these associations to several symptom domains and identified transcriptional signatures associated with distinct functional pathways, including GABAergic and glutamatergic neurotransmission, metabolic processes, and intracellular signal transduction, across brain regions associated with distinct symptomatic profiles in a sex-specific fashion. In most cases, these associations were specific to males or to females with MDD, although a subset of gene modules associated with common symptomatic features in both sexes was also identified. Together, our findings suggest that the expression of distinct MDD symptom domains is associated with sex-specific transcriptional structures across brain regions.

## Introduction

Major depressive disorder (MDD) is a highly pervasive and recurrent disease affecting yearly more than 280 million people worldwide^1^. Despite its major burden on modern societies, current strategies to treat the syndrome remain relatively inefficient with only a third of patients showing complete remission and roughly two third exhibiting various levels of recurrence^2^. These limitations in MDD treatment likely result in large part from its clinical heterogeneity.

Based on clinical symptoms, MDD is a highly heterogeneous syndrome defined by the expression of depressed mood and anhedonia^3^. These two core symptoms are accompanied variably by cognitive impairment, anxiety, weight change, fatigue, agitation, sleep abnormalities, feelings of worthlessness, recurrent thoughts of death and suicidal ideations, among other symptoms^3^. These symptoms not only vary across patients but their expression also evolves in individuals along with the chronicity of the illness. Accordingly, previous attempts to cluster and treat patients based on their symptomatic profiles have shown only modest success^4^. Part of this lack of success can be explained by the way the syndrome is defined and diagnosed. Indeed, as of now, MDD diagnosis is based entirely on standardized, but yet subjective, behavioral measures, and the severity of depressive episodes is obtained by summing up symptoms’ presence instead of their intensity^4^. However, these symptoms are broadly different in many dimensions, including their underlying biological substrates and associated molecular mechanisms^5,6^.

Findings from functional imaging studies have culminated on the creation of different models linking the expression of specific MDD-relevant clinical features with the activity of distinct brain regions and circuits^7,8^. For instance, hyper-connectivity/activation between the default mode network—Negative Affect Sad and Negative Affect Threat networks—and their respective brain regions, has been associated with the expression of rumination^9^, sadness and hopelessness (negative bias)^9,10^ and threat dysregulation (scariness and sense of failure)^11,12^, respectively. These studies also suggest that the expression of anxiety^13,14^, inattention and cognitive dyscontrol (poor concentration, indecisiveness)^4,15–17^ and anhedonia^18,19^ associates with global hypo-connectivity/activation of the Salience, Attention and Cognitive Control and Positive Affect Happy networks, respectively. These findings support the idea that changes in the activity of specific brain regions and circuits drive the expression of distinct clinical features of MDD, even though the molecular mechanisms underlying these functional changes in each respective brain region remain poorly understood.

Transcriptional changes affecting not only gene expression but also the organization of gene networks have been reported across several brain regions and circuits in post-mortem brain tissues of MDD patients and mouse chronic stress models^20–28^. For instance, detailed analyses revealed the role of gene networks in mediating stress susceptibility in a sex-specific fashion by interfering with intracellular cascades regulating neuronal activity^24,25^. More recently, transcriptional signatures in brains of MDD patients have been associated with trait versus state depression, revealing gene profiles associated with the transition between both clinical states in males^29^. However, none of these studies have been able to show whether any of these transcriptional signatures associate with the expression of the symptomatic manifestations of MDD in males and females.

Data driven system-based approaches have shown their advantages over conventional methods in revealing pathogenic etiologies for complex and heterogeneous neuropsychiatric disorders. Network-based analyses in particular provide the tools and statistical approaches to classify sub-types of complex diseases according to their molecular profiles. These methods have been used to highlight the molecular architecture underlying the expression of several complex neuropsychiatric disorders such as Alzheimer disease, autism, bipolar disorder, schizophrenia and MDD^25,30–32^. Here, we used network analyses to evaluate the potential association between transcriptional signatures across brain regions and the expression of distinct symptom domains relevant to MDD in males and females.

## Methods

Brain tissues were obtained from the Douglas Bell Canada Brain Bank (Douglas Mental Health Institute, Verdun, Québec and from the University of Texas Southwestern Medical Center Brain Bank. In total, analyses were performed on 89 samples including 25 male MDD, 25 female MDD, 17 male CTRL (healthy controls) and 22 female CTRL. Sociodemographic and clinical information including sex, phenotype (MDD, CTRL), age, pH, postmortem interval (PMI), treatment history, smoking history, history of early life adversity, cause of death, presence of drug and/or alcohol abuse and cohort (Montreal, Texas) is listed in **Suppl. Table 1**. All analyzes were performed on six brain regions including the anterior insula (aINS), orbitofrontal cortex (OFC; BA11), cingulate gyrus 25 (BA25; cg25; vmPFC [ventromedial prefrontal cortex]), dorsolateral PFC (BA8/9; dlPFC), nucleus accumbens (NAc) and ventral subiculum (vSub). Overall, we sequenced RNA from 41 new human brains and combined this dataset with RNA profiles from 48 brains published by us before^28^. Postmortem tissue from all six brain regions was carefully dissected at 4°C after having been flash-frozen in isopentane at −80°C. All dissections were performed by histopathologists using reference neuroanatomical maps^33^.

Psychiatric history and socio-demographic information was obtained via psychological autopsies carried out by trained clinicians using the same methods in case and control groups^34,35^. Diagnosis and clinical information including symptomatic profiles were obtained using DSM-IV criteria by means of SCID-I interview adapted for psychological autopsies^3,36^. Nine main categories of symptoms were recorded, including depressed mood, loss of interest or pleasure, change in appetite/weight, insomnia/hypersomnia, psychomotor agitation/retardation, fatigue or loss of energy, low self-esteem, difficulty in concentration/indecision and recurrent suicidal thoughts. Notably, since depressed mood, anhedonia, fatigue and recurrent suicidal thoughts were expressed by every MDD patient, we did not include those symptoms in our analysis (**Suppl. Table 2**)

### RNA sequencing

RNA from human postmortem brain samples was extracted using the RNeasy micro kit with Trizol, followed by DNase I treatment, as described by the manufacturer (Qiagen). RNA integrity (RIN) and concentration was quantified using a Bioanalyzer (Agilent). RNA libraries were synthesized from 1 μg of RNA using the ScriptSeq Complete Gold Kit (Epicentre, Illumina) including an initial ribosomal RNA depletion step. Each library was spiked with an external RNA sample as a control as suggested by the manufacturer (Thermofisher). Samples were barcoded and sequenced in multiplex (8 per lane) twice at a depth of 50 million reads (50 bp paired-end) per sample on Illumina HiSeq2500.

### Data processing

Sequencing data from all 89 samples for all 6 brain regions were analyzed using the same criteria. Sequencing quality and trim reads were assessed using FASTQ and FASTX-toolkit. TopHat was used to align paired-end reads to the GENCODE 2019 (GRCh38.p12) human annotation. Overall, every sample included in this study passed QC assessment. Reads for every sample were counted using HTSeq. A gene was considered the union of all its exons in any known isoforms, based on GENCODE annotation. Any reads that fell in multiple genes were excluded from the analysis. Threshold for filtering out genes expressed at low levels was set to <5 reads in at least 20% of the samples per group as described previously^28^.

We adapted multiple preprocessing steps to ensure both statistical and biological relevance. Gene expression was first transformed and normalized using the *voom* function in the Limma package^37,38^. Batch effect and potential unwanted sources of variance in gene expression across all samples was identified through RUVseq using spike-in controls^39^. As expected, the effect of batch (new and previous cohort) was found to be significant for every brain region. The top first factor was extracted and included as a covariate in the differential expression analysis. We then performed a principal component analysis (PCA) to reveal the effect of clinical and technical covariates on variations of gene expression. We identified significant effects for PMI, pH, cohort, drug abuse and RIN in the aINS; age, PMI, pH, childhood abuse, cohort, drug abuse and RIN in the OFC; age, pH, cohort, drug abuse and RIN in the vmPFC; age, PMI, pH, cohort, drug abuse and RIN in the dlPFC; age, PMI, childhood abuse, cohort, drug abuse and smoking in the NAc; and PMI, cohort, drug abuse and RIN in the vSub (**Suppl. Table 3**). The effects of these covariates were adjusted in our downstream differential expression and gene co-expression network analyses.

#### Differential expression analysis

Differentially expressed genes (DEGs) were identified through a generalized linear model (GLM) implemented in limma with sex (male and female) and phenotype (MDD and CTRL) as main factors. A single GLM was performed for each brain region controlling for every region-specific covariate identified through our PCA and RUV analyses. An individual gene was called differentially expressed if the nominal *P*-value of its *t*-statistic was ≤0.05. Globally, this approach allowed the identification of genes differentially expressed in males and females with MDD while controlling for baseline variations in gene expression along with the effects of clinical and technical covariates across every brain region.

#### Transcriptional overlap analysis

We used a rank-rank hypergeometric overlap (RRHO) analysis^40,41^ to measure transcriptional overlap between males and females with MDD. RRHO was also used to confirm the reproducibility of our results by overlapping results from the current analysis with previously published transcriptional maps in males and females with MDD^25^. Gene lists were ranked and signed according to their degree of differential expression in male MDD and female MDD versus CTRL, respectively (–log_10_(*P*-value)) multiplied by sign of the *t*-statistic. For each comparison, a matrix of hypergeometric *P*-values was created as a result of the iterative statistical tests evaluating the proportion of ranked genes differing from one condition to another. Multiple testing correction was performed using the Benjamini and Yekutieli (BY) method^42^. Adjusted *P*-values were finally heat-mapped with each pixel representing the adjusted –log_10_ hypergeometric *P*-values of the transcriptional overlap between males and females with MDD.

#### Gene ontology analysis

Gene ontology analysis was performed using g:Profiler2^43^ on DEG lists from males and females with MDD across all brain regions with significant enrichment fixed at FDR<0.05.

### Gene co-expression network analysis

Weighted gene co-expression network analysis (WGCNA)^44^ was used as a systems biology approach to identify modules of highly co-expressed genes. Multiple iterations of WGCNA were performed according to our objectives including (1) for males and females with and without MDD for every brain region (24 networks) and (2) combining males and females with and without MDD for every brain region (6 networks). Five samples were detected as outliers before running network analysis. These outliers were identified via the construction of an Euclidean distance-based sample network with standardized connectivity <-3.5 as the exclusion connectivity threshold. These outliers were removed from the final network construction. Network construction was adjusted for the same covariates used in the differential expression analysis (**Suppl. Table 3**). Weighted gene co-expression networks were built with a matrix of biweight midcorrelation between all gene pairs which was converted to an unsigned adjacency matrix using a soft threshold power and then transformed into a topological overlap matrix for modular structure detection^45,46^. Highly co-regulated genes were identified through average linkage hierarchical clustering to create groups of genes, with a subsequent dynamic tree cut to explore clusters in a nested dendrogram, with each branch corresponding to a module. Each module was named by a unique arbitrary color and associated with an ontological term using g:Profiler2 from Bioconductor. FDR *P*-values and fold enrichment for each module were reported. Genes with the highest intramodular connectivity (top 5% highest intramodular connectivity) were considered as hub genes. Network organization was represented through Cytoscape v3.9.1^47,48^.

Modular enrichment for DEGs was assessed using the GeneOverlap package^49^ from Bioconductor. Enrichment was tested for genes significantly upregulated or downregulated in each module in males and females across all 6 brain regions. Fisher’s exact tests were used to perform enrichment assessment with significance fixed at P_adj_<0.05.

#### Module differential connectivity

We used module differential connectivity (MDC) to quantify differences in co-expression network organization between male MDD and female MDD compared to controls and male MDD versus female MDD across brain regions. MDC is determined by calculating the ratio of connectivity between all gene pairs in a module in one condition (phenotype, sex or brain region) to that of the same gene pair in another condition. MDC values larger than 1 indicate gain of connectivity (GOC) or stronger co-expression between genes, while values lower than 1 indicate loss of connectivity (LOC) or weaker co-expression between genes. The statistical significance of MDC was adjusted for multiple testing using the FDR permutation method^50^.

#### Module preservation

Module preservation was carried out to assess whether gene modules in males and females were preserved in the opposite sex, respectively and across brain regions in males and females. Module preservation was computed with the preservation statistics of the WGCNA package. Network preservation statistics do not require independent module identification in a test group. The approach evaluates the preservation of connectivity patterns of the member genes and the distinctiveness of a module as a whole from other modules. Module preservation can be established by four complementary statistics including median rank, *Z*_density_, *Z*_connectivity_ and *Z*_summary_. *Z*_density_ and *Z*_connectivity_ statistics are the standardized preservation statistics for density and connectivity, respectively, while *Z*_summary_ is the average of *Z*_density_ and *Z*_connectivity_. Preservation in this study was established through the Z_summary_ measures. Modules with a Z_summary_ score higher than 10 were considered preserved, as recommended^51^.

#### Clinical association with gene expression and transcriptional modular structure

Association between gene expression and clinical symptoms across brain regions was calculated by means of hierarchical clustering with complete linkage, using Pearson correlation coefficient, on the 200 strongest DEGs in each brain region. Heatmaps were made using CPM for the top 200 DEGs versus samples with and without the expression of each symptom in males and females separately.

Symptom association with network structure was performed by calculating point-biserial correlation coefficients between clinical symptoms (dichotomous variable) and module eigengene values (continuous variable). Module eigengene values are defined as the first principal component of each module. It represents global variance within each module and is calculated using the *moduleEigengenes* function in WGNCA^45^. *P*-values were adjusted for multiple testing using permutation test (1000 permutations). Finally, relationship between gene membership and symptomatic expression was assessed by means of correlation between the gene significance (GS) and module membership. This allowed identifying whether specific genes in each module contribute to the module’s association with the clinical symptoms. Module enrichment for genes associated with each symptom was measured via Fisher’s exact test with *P*-values corrected for multiple testing using Benjamini-Hochberg^52^.

### Statistical analysis

Although sample size calculation was not performed, the sample size in this study is justified based on several previously published reports using similar or even smaller sample sizes and showing the power to detect significant statistical differences. RNA-seq gene expression data for differential expression was normalized. In total, 89 samples including 25 male MDD, 25 female MDD, 17 male CTRL and 22 female CTRL, from six brain regions (total 534 samples) were included in this study. Overall, transcriptional profiles were generated for 41samples in six brain regions and combined with RNA profiles from 48 samples published by us before^28^. Differential expression was not corrected for multiple testing. RRHO analysis and FET were corrected using the Benjamini-Hochberg method. Network analysis included network construction, module differential connectivity, GO enrichment, module preservation and module differential expression enrichment, all of which were corrected for multiple testing. Association between clinical symptoms and modular structures in males and females with MDD were calculated with point-biserial correlations and correlations between the gene significance (GS) and module membership. *P*-values calculated for these coefficients were adjusted using permutations and Benjamini-Hochberg adjustment. Details of each analysis are provided above in each respective section.

## Results

The main objective of this study was to determine whether transcriptional signatures across different brain regions associate with the expression of specific symptomatic profiles in males and females with MDD. To do so, we first mapped transcriptional signatures in males (n=25) and females (n=25) with MDD compared to healthy control subjects (17 males and 22 females) across six brain regions (see **Suppl. Table 4** for detailed cohort composition by brain region). We then established the degree of transcriptional changes across six brain regions in males and females with MDD using differential expression analysis and WGCNA. We explored male and female transcriptional profiles to identify unique and shared associations with the expression of specific symptoms of MDD in both sexes. Finally, we identified hub and node genes for each network and calculated their contribution to the association between their respective module and specific symptoms of MDD in both sexes. Overall, our results show that specific transcriptional signatures associate with the expression of distinct symptomatic profiles and reveal the molecular substrates underlying the expression of MDD and its clinical manifestations across the brain of males and females.

### Differential expression reveals sex- and brain region-specific transcriptional signatures in MDD

We first used differential expression analysis to identify genes significantly up- or downregulated across six brain regions of males and females with MDD. Our analyses revealed a large number of genes differentially expressed (DEG, p<0.05) in males and females across every brain region with a small proportion of overlap between the two sexes (**Fig. 1a; Suppl. Tables 5-6**). In total, we identified between 3.2% up to 35.9% of overlapping DEGs in males and females with MDD across brain regions. The overlap was smaller in the NAc (3.2%) and vSub (4%) between males and females with MDD, while greater overlap was seen in PFC regions: 35.9% in dlPFC and 11.9% in vmPFC.

**Figure 1.**
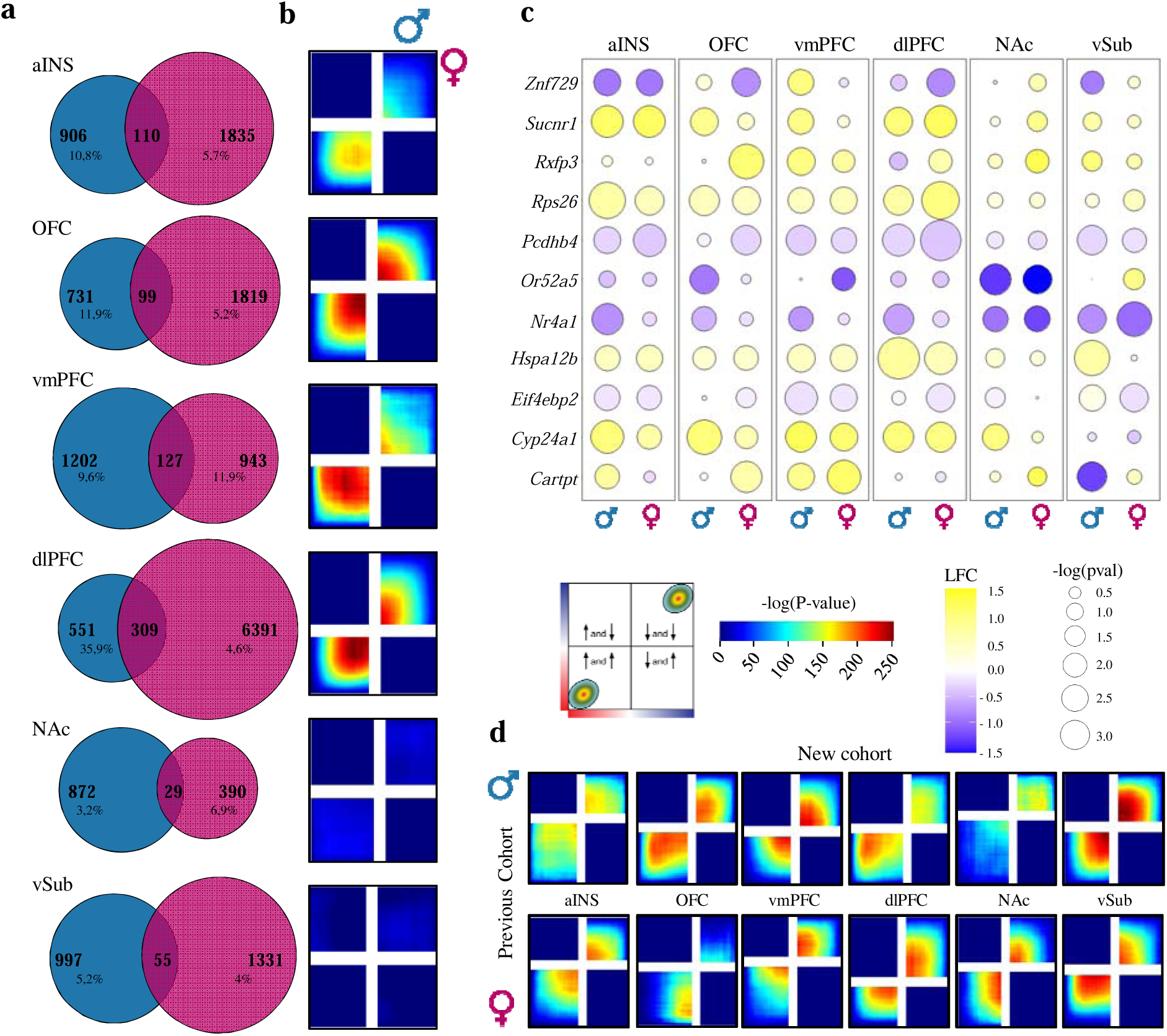
Differential gene expression patterns differ in males and females with MDD, revealing sex- and brain region-specific transcriptional profiles. (**a**) Venn diagrams of DEGs (P<0.05) showing low overlap between male MDD (blue) and female MDD (pink) across brain regions. (**b**) RRHO maps displaying transcriptional overlaps between male MDD and female MDD across brain regions. Signals in the bottom left and upper right quadrants represent an overlap for commonly upregulated and commonly downregulated genes, respectively. The color bar represents the degree of significant (-log (P_adj_-value) overlap between transcriptional signatures in males and females with MDD. (**c**) Bubble plot showing the most variable genes differentially expressed in male MDD and female MDD across six brain regions. Colors represent log fold change values, with blue for genes downregulated and yellow for genes upregulated in MDD versus control conditions. The radius of the circles shows significance levels according to their *P*-values. (**d**) RRHO maps representing transcriptional overlaps between the new cohort sequenced in this paper and a previously published cohort^27^.

To further characterize the transcriptional overlap between males and females with MDD, we used a rank-rank hypergeometric overlap (RRHO) analysis to compare transcriptional signatures from both sexes without restricting our analysis to stringent statistical thresholds^40,41^. Interestingly, our results revealed a significant overlap (max. *P*-value=1.0E-250) for genes commonly up- or downregulated in both males and females, but only in cortical regions including the OFC, vmPFC dlPFC and to a lower extent the aINS (**Fig. 1b**). In contrast, RRHO analysis revealed a lack of transcriptional overlap in limbic structures including the NAc and vSub (**Fig. 1b**). Importantly, our analysis at the gene level supports these observations with the most variable genes from our datasets showing sex- and brain region-specific transcriptional changes (**Fig. 1c**). For instance, *Znf729*, *Rxfp3*, *Or52a5*, *Eif4ebp2* and *Cartpt* exhibit opposite transcriptional patterns between males and females with MDD across brain regions. Furthermore, genes such as *Nr4a1*, *Hspa12b*, *Pcdhb4*, *Rps26* and *Sucnr1* show consistent changes across brain regions while others exhibit region-specific transcriptional regulation.

We then used gene ontology analysis to identify functional features enriched with DEGs across each brain regions. As expected, this analysis identified functional terms broadly different between males and females across each brain region (**Suppl. Tables 7-8**). However, we also identified a subset of functional domains shared between males and females with MDD, including *GABAergic synaptic function* in the aINS, *neuropeptide signaling pathway* in the vmPFC and *AMPA receptor activity* and *neurotransmitter receptor complex* in the vSub (**Suppl Fig. 1**). These findings suggest that sex-specific transcriptional changes converge onto some similar functional alterations across brain regions in males and females.

Finally, to confirm the reproducibility of our findings, we overlapped our transcriptional profiles with those from our previous work^28^ using DEG and RRHO analyses. Importantly, our examination revealed a strong and consistent overlap between results obtained in this new cohort and those from males and females with MDD previously published on the same brain regions tested both at the gene (DEG) and broader (RRHO) signature levels (**Suppl Fig. 2; Fig. 1d**). Overall, results from our analyses are consistent with previous comparisons of DEG profiles in males and females with MDD^22,27,28^ and suggest that transcriptional signatures in both sexes exhibit broad differences but with a certain level of overlap, mainly in cortical brain regions, ultimately affecting genes converging onto common functional pathways across brain regions in males and females with MDD.

**Figure 2.**
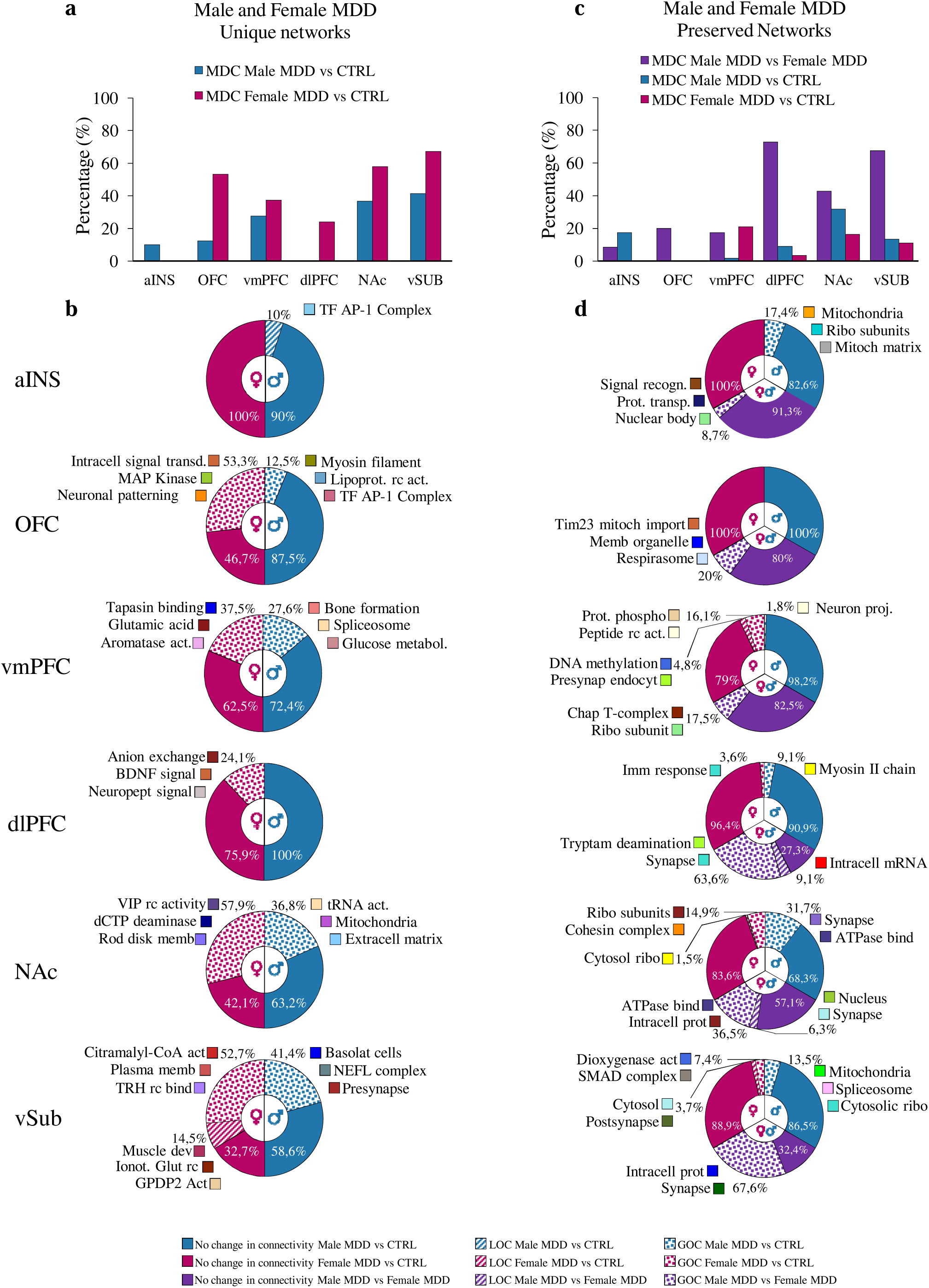
Changes in the structural connectivity of sexually preserved/unique modules contribute to the expression of MDD. **(a)** Proportion of unique modules with significant MDC in MDD versus CTRL in males (blue) and females (pink). **(b)** Proportion of unique modules with significant LOC (striped area) and GOC (dotted area) and their functional ontological terms. **(c)** Proportion of preserved modules with significant MDC in MDD versus control in males (blue) and females (pink) and in male MDD versus Female MDD (purple). **(d)** Proportion of preserved modules with significant LOC (striped area) and GOC (dotted area) and their functional ontological terms.

### WGCNA highlights region-specific gene networks associated with MDD in males and females

We next used WGCNA to construct the transcriptional organization of gene networks in males and females with and without MDD across all six brain regions. Importantly, as opposed to previous strategies of pooling brain regions^28^, the size of our current combined cohort allowed us to create sex-specific gene modules for all six brain regions investigated. Gene ontology analysis was used to associate a functional term to each of these modules. In total, we identified between 20 to 109 gene modules composed of between 50 to 7662 genes across every brain region representing in an unbiased manner the vast majority of functional domains relevant to brain activity (i.e., *synaptic function*, *metabolic function*, *cytoskeletal plasticity*, *immune function*, etc.) in males and females with and without MDD (**Suppl. Fig. 3**; **Suppl. Table 9**).

We started by testing the extent to which the transcriptional organization of gene networks is preserved across the brain of males and females with MDD. To do so, we calculated Z_summary_ values for modules in the male MDD group and considered every module with a Z_summary_ score higher than 10 as being preserved in the female MDD. Not surprisingly, we found a significant proportion (from 35% to 77%) of male gene networks preserved in females across all six brain regions (**Suppl. Fig. 4**). The NAc (76.8%), aINS (69.7%) and vmPFC (66.3%) showed the highest level of preservation in modules found in male versus female MDD, while a smaller but still considerable proportion of gene networks were preserved in the vSub (34.6%), OFC (52.2%) and dlPFC (55.0%). Together, this suggests that similar transcriptional structures are involved in the control of fundamental cellular processes across brain regions in males and females.

We then tested whether these modules, either preserved or unique in both sexes, were associated with changes in intramodular connectivity. Intramodular connectivity represents the strength of the overall correlation values amongst genes from the same module. Change in intramodular connectivity between two conditions (i.e., MDD versus CTRL), defined as module differential connectivity (MDC), has been associated with dysfunctional organization of transcriptional structures in stressed mice and humans with MDD^25,28,53,54^. Interestingly, our analysis suggests that a large proportion of unique modules in males and females with MDD associate with a significant MDC compared to their respective controls. In female MDD, this proportion reaches 67.3% in vSub, 57.9% in the NAc and 53.3% in the OFC, while no unique module in female aINS showed differential connectivity (**Fig. 2a, b**). Similar, although lower, proportions were found in males with MDD: the vSub (41.4%), NAc (36.8%) and vmPFC (27.9%) showed the highest proportion of modules associated with intramodular changes in structural connectivity compared to their respective controls (**Fig. 2a, b**). For both males and females, modules with a significant GOC or LOC were enriched for genes relevant for different functional terms, such as *transcription factor activity* (Male aINS and OFC), *BDNF signaling* (female dlPFC), *neuropeptide activity* (female NAc) and *synaptic activity* (male vSub) in a region-specific fashion (**Fig. 2b**).

In contrast, a much smaller proportion of modules preserved between males and females with MDD exhibit significant MDC compared to their respective control conditions. Indeed, with the exception of the NAc (31.7%) in male MDD and the vmPFC (21%) in female MDD, the proportion of sex-preserved modules associated with a significant MDC compared to their respective controls was lower than 20% in every brain region (**Fig. 2c, d**), with the OFC and aINS showing no module associated with MDC in either males or females with MDD. On the contrary, our analysis revealed a large proportion of modules preserved in males and females with MDD associated with a significant MDC when compared to the other sex (Male MDD versus Female MDD). Indeed, this proportion reached 72.7% in the dlPFC, 67.6% in the vSub and 42.9% in the NAc, while lower levels were found in the aINS (8.7%), vmPFC (17.5%) and OFC (20.0%) (**Fig. 2c, d**). Modules preserved in both sexes associated with a significant GOC or LOC were also enriched for genes relevant for different functional terms, including *synaptic function* (aINS, vmPFC, dlPFC, NAc, vSub), *function of the mitochondria* (aINS, OFC, vSub), *intracellular protein signaling* (aINS, vmPFC, NAc, vSub) and *nuclear control of gene expression* (aINS, vmPFC) (**Fig. 2d**). Together, this suggests that a significant proportion of the transcriptional organization of gene networks is shared across the brain of males and females with MDD. Nevertheless, despite this high level of homology, our findings demonstrate that both unique and sexually-preserved gene modules contribute to the expression of MDD distinctly in males and females via changes in their structural connectivity and underlying biological functions. Furthermore, brain regions such as NAc and dlPFC are more importantly associated with MDD in both sexes than the aINS.

### Transcriptional associations with symptomatic features in males and females with MDD

Our analyses to this point show that networks of co-expressed genes in males and females contribute to the expression of MDD in both sexes. We next tested whether transcriptional signatures across brain regions in males and females with MDD associate with specific symptomatic profiles in both sexes. Clinical information for each sample was obtained by means of post-mortem psychological autopsies as described before^34,35^. Globally, symptomatic data obtained through this approach provided information on change in appetite/weight, insomnia/hypersomnia, psychomotor agitation/retardation, low self-esteem and difficulty in concentration/indecision (**Suppl. Table 2**) with similar proportions in males and females with MDD. We first ran a hierarchical clustering analysis to test whether the expression of specific symptomatic features associates with variations in gene expression across brain regions in males and females. Interestingly, our analysis in females revealed clear patterns of gene expression across all brain regions in samples expressing insomnia/hypersomnia (**Fig. 3a**). Similar patterns were observed in the aINS, vmPFC and dlPFC for samples with psychomotor agitation/retardation and for changes in appetite/weight in the vmPFC and vSUB. In contrast, no such clear patterns were identified in males with MDD (**Fig. 3a**).

**Figure 3.**
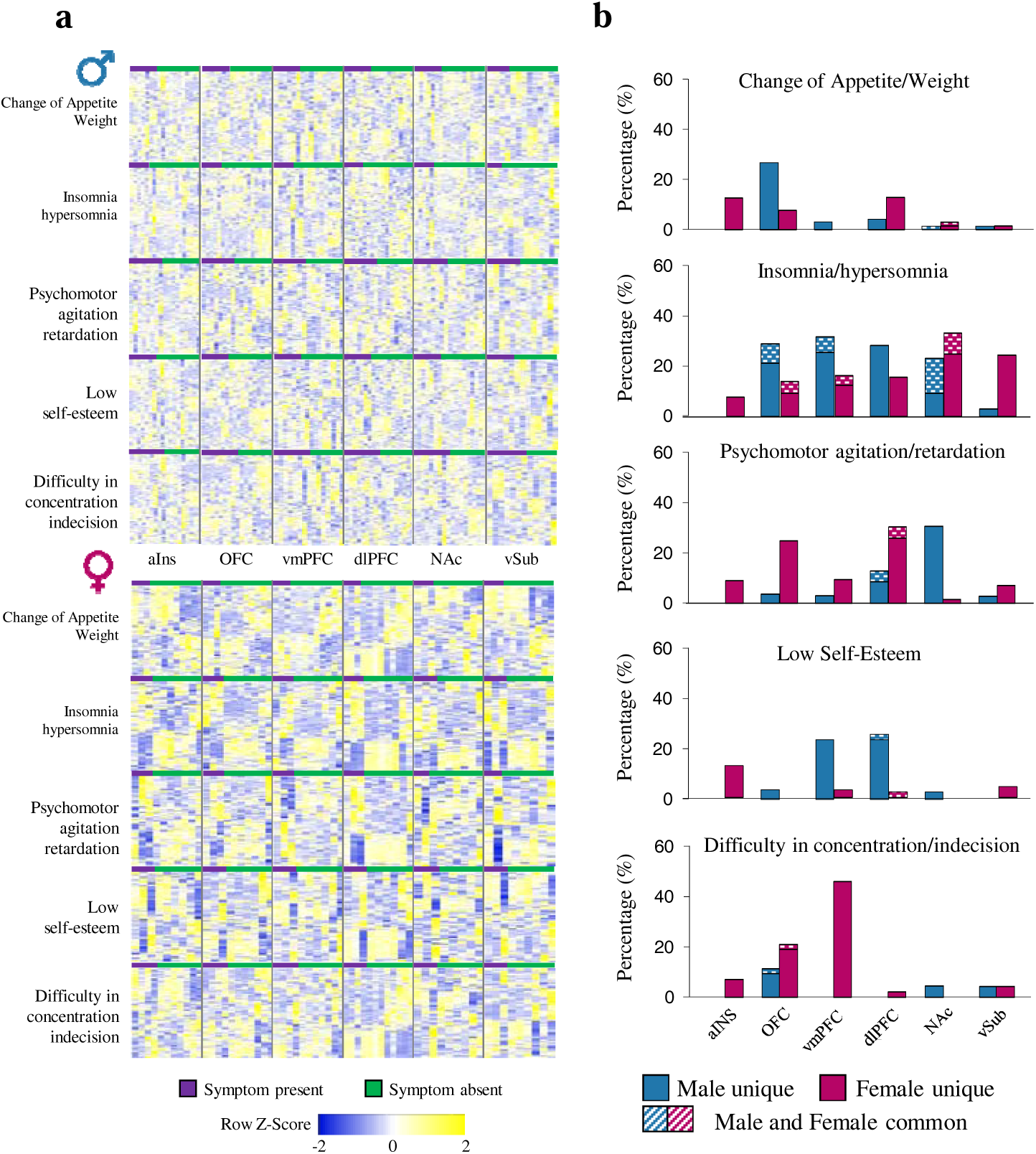
Symptomatic profiles in MDD associate with transcriptional alterations across brain regions in males and females. **(a)** Heatmaps displaying hierarchical clustering of the most variable genes according to the presence (blue) or absence (orange) of symptoms. **(b)** Proportion of gene co-expression modules associated with symptomatic features in males (blue) and females (pink) with MDD. Stippled areas show percentage of modules in males and females associated with the same symptoms in both sexes.

We then expanded our analysis to assess whether the integrative and correlative features of WGCNA would provide further advantages in revealing significant associations between male and female MDD symptomatic profiles and transcriptional gene networks. To do this, we constructed gene networks combining transcriptional profiles from males and females with and without MDD (**Suppl. Fig. 5**) and calculated module eigengene values for samples in every brain region by extracting the first principal component from each identified module. For every given symptomatic feature, we measured the associations between that specific symptom and the module eigengene values using biserial correlations. This approach identified several modules associated with the expression of each symptom of MDD, although these associations differed by sex and brain region (**Fig. 3b**). For instance, the largest proportion of modules in males associated with change of appetite/weight was found in the OFC (26.9%), psychomotor agitation/retardation in the NAc (31.0%), low self-esteem in the dlPFC (26.1%) and difficulty in concentration/indecision in the OFC (11.5%). In females, the highest proportion of modules associated with change of appetite/weight was found in the dlPFC and aINS (13.0% and 12.7%, respectively), psychomotor agitation/retardation in the dlPFC (30.4%), low self-esteem in the aINS (12.7%) and difficulty in concentration/indecision in the vmPFC (46.0%). Overall, the expression of insomnia/hypersomnia was associated with a larger proportion of gene networks across the brain, in both males and females, compared to other symptoms (**Fig. 3b; Suppl. Fig. 6**).

We next investigated which of these gene networks are most strongly associated with the expression of specific symptoms across brain regions of males and females with MDD. For instance, *Ivory* in the OFC (**Fig. 4a**) is the gene network most strongly associated with the expression of change in appetite/weight in males with MDD (r=0,76; p_adj_<0,001) (**Fig. 4b**). Noticeably, this network is also associated with the expression of insomnia/hypersomnia (r=0,68; p_adj_<0,005) (**Fig. 4c**) and difficulty in concentration/indecision (r=0,73; p_adj_<0,005) (**Fig 4d**) in male MDD. *Ivory* is enriched for genes associated with synaptic signaling, most importantly GABAergic neurotransmission (p_adj_<5.0^E-8^) (**Fig. 4f**). Alterations of the GABAergic system in the PFC have been frequently associated with the expression of MDD^55,56^, although its role in specific symptomatic features has never been reported. Interestingly, *Ivory* is depleted of DEGs but is enriched for genes significantly associated with the expression of each three symptoms in males (Fisher Exact Test (FET): change in appetite/weight, p<1.0^E-18^; insomnia/hypersomnia, p<5.0^E-11^; difficulty in concentration/indecision p<5.0^E-17^) (**Fig. 4e**), including all three hub genes, namely, *Gad1* and *Gad2* which encode glutamate decarboxylase 1 and 2 and *Nxph1* which encodes neurexophilin 1 (**Fig. 4a**). In addition to *Ivory* in the OFC, we also identified *Darkviolet* in the NAc and *Darkred* in the dlPFC which are associated with the expression of psychomotor agitation/retardation (r=-0,69; p_adj_<0,0001) and low self-esteem (r=-0,83; p_adj_<0,0001) in males with MDD, respectively (**Suppl. Fig. 7**).

**Figure 4.**
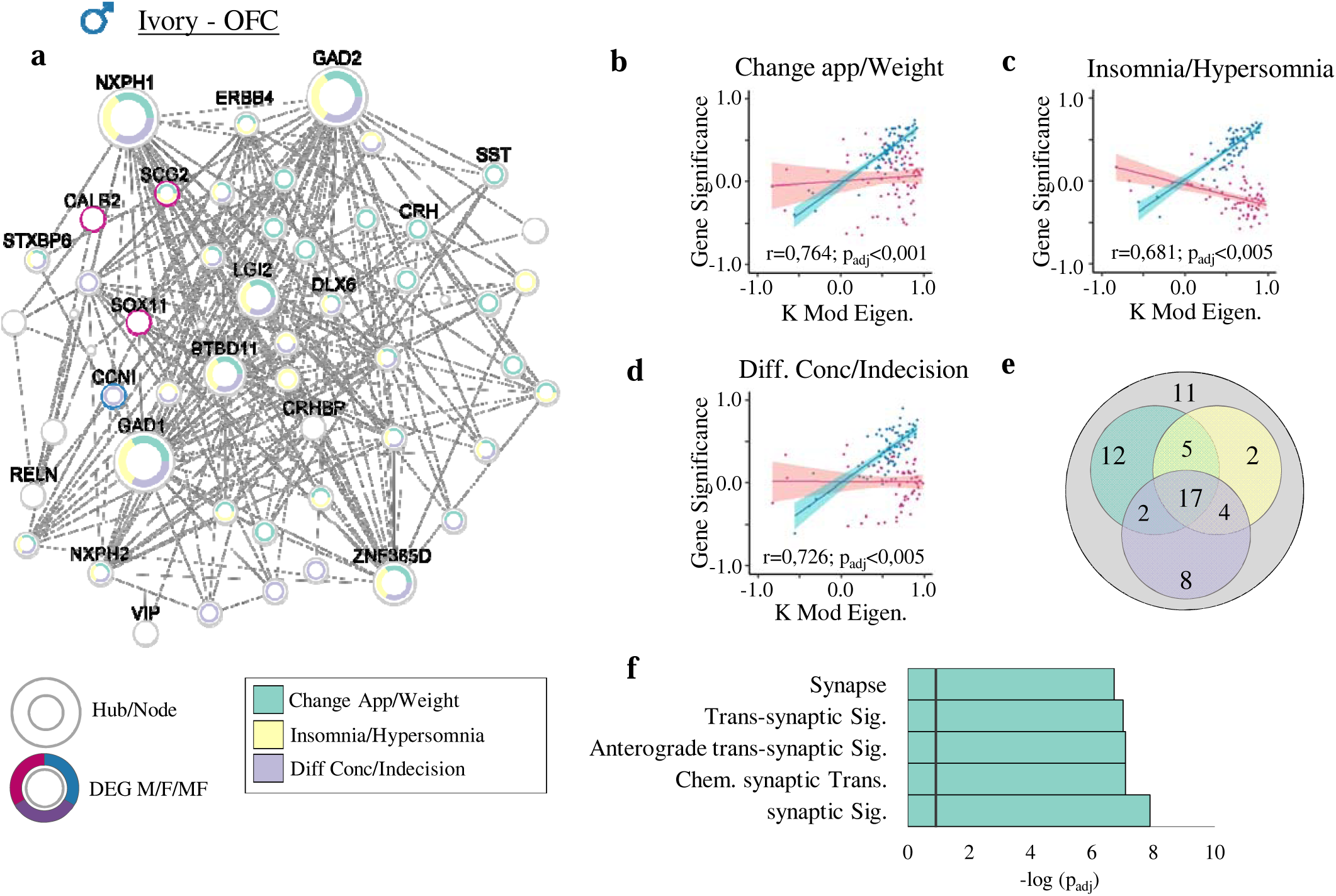
Most significant module (*Ivory*) associated with the expression of change in appetite/weight, insomnia/hypersomnia and difficulty in concentration/indecision detected in male OFC. **(a)** Hubs and nodes representation of *Ivory* and their associations with clinical symptoms of MDD. The inner colors show with which symptom each gene (hub/node) is associated, with green for change in appetite/weight, yellow for insomnia/hypersomnia, and purple for difficulty in concentration/indecision. The surrounding color shows if the gene is differentially expressed in male (blue), female (pink) or both (violet). Hubs and nodes are defined by the size of the circles. **(b-e)** Correlation between module membership (KME) and gene significance values of the association between each gene and relevant symptomatic feature, for genes co-expressed in *Ivory,* in males (blue) and females (pink) with the 95% confidence interval band. € Venn diagram showing the number of genes in the *Ivory* module associated with at least one out of the three symptoms, with change in appetite/weight in green, insomnia/hypersomnia in yellow and difficulty in concentration/indecision in purple. **(f)** GO enrichment for the *Ivory* module for the first five most significantly enriched GO terms. The black vertical line indicates the significance threshold (p_adj_ <0,05).

In females, our analysis pointed to *Darkorange* in the aINS as a module associated with the expression of change in appetite/weight uniquely in female MDD (**Fig. 5a**). *Darkorange* is enriched in genes relevant to synapse (p_adj_<1.0^E-9^) and cell junction (p_adj_<1.0^E-8^). Importantly, *Darkorange* in female MDD is enriched in downregulated DEGs (p_adj_<3.0^E-6^) and also associated with the expression of change in appetite/weight (p_adj_<1.0^E-57^). In fact, 39% of all genes in *Darkorange* are associated with the expression of this symptom in female MDD (p_adj_<5.0^E-26^), including the seven hub genes among which *Clstn1* and *Clstn3* which encode transmembrane protein calsyntenin family members and *Pi4ka* which encodes phosphatidylinositol (PI) 4-kinase were also significantly downregulated in the aINS of females with MDD (**Fig. 5a**). As well, we identified *Saddlebrown* in the dlPFC (**Fig. 5e**) to be significantly associated with the expression of psychomotor agitation/retardation (r=0,891; p_adj_<0,0001) (**Fig. 5f**) in female MDD. This module (cellular protein catalytic process; p_adj_<0.0005) (**Fig. 5h**) is strongly enriched for upregulated DEGs (p_adj_<5.0^E-37^) (**Fig. 5f**) and with the expression of psychomotor agitation/retardation in the dlPFC of female MDD (34.1%; p_adj_<1.0^E-17^) (**Fig. 5g**), including all 5 top hub genes *Selenot*, *Actr3*, *Chmp2b*, *Sgpp1* and *Tm9sf3*. Additional modules strongly associated with the expression of insomnia/hypersomnia (*Skyblue*-OFC; r=-0,747; p_adj_<0,0001), low self-esteem (*Purple*-aINS; r=-0,827; p_adj_<0,0001) and difficulty in concentration/indecision (Palevioletred3-vmPFC; r=-0,60; p_adj_<0,0001) in females with MDD are highlighted in **Suppl. Fig. 8**.

**Figure 5.**
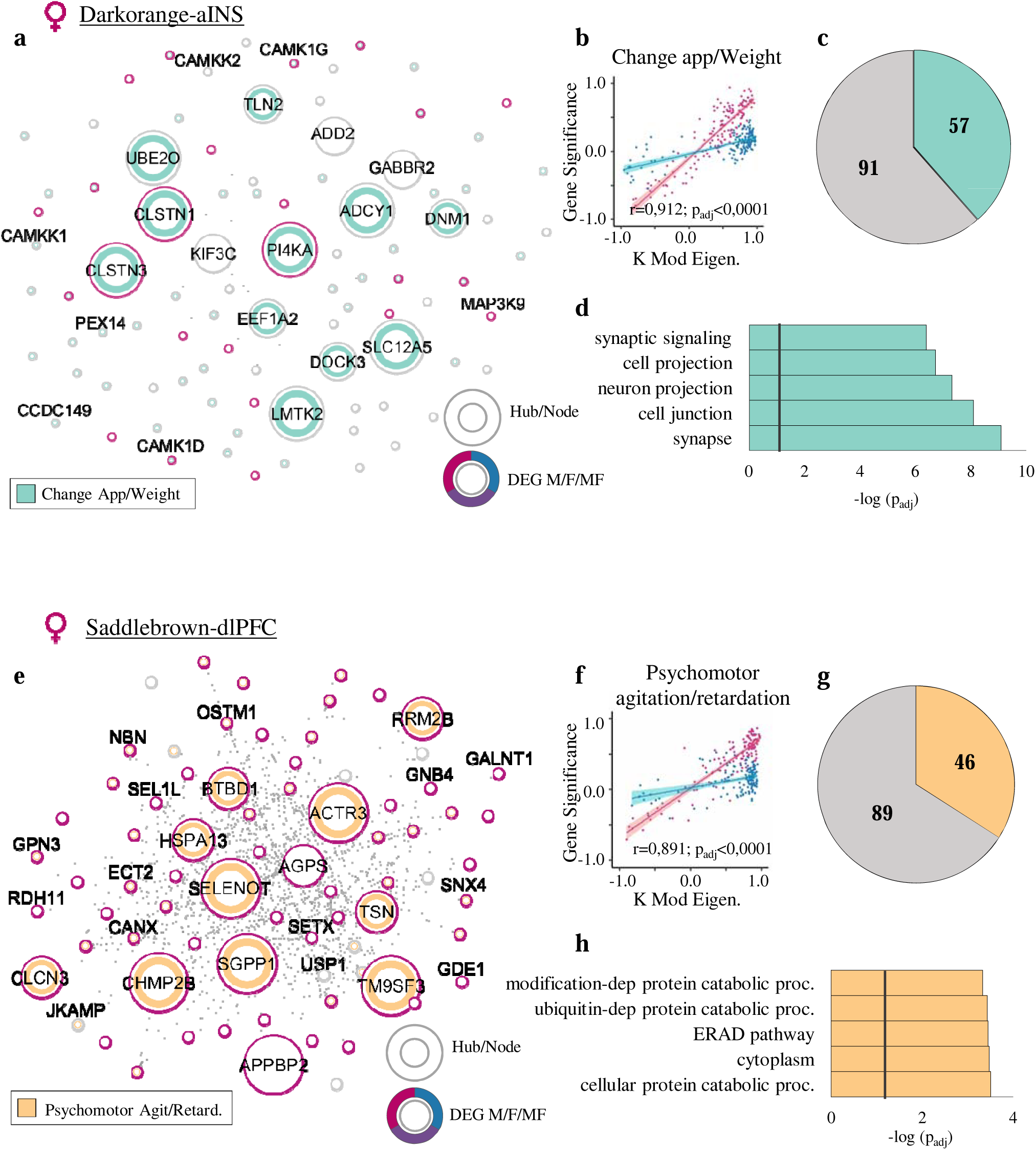
The most significant modules (*Darkorange* and *Saddlebrown*) associated with the expression of change in appetite/weight and psychomotor agitation/retardation in female MDD detected in the aINS and dlPFC, respectively. **(a,e)** Hubs and nodes representation of **(a)** *Darkorange* and **(e)** *Saddlebrown* associated with their respective clinical symptoms. **(a)** The inner color in green shows the genes significantly associated with the expression of the symptom: change in the appetite/weight. **(e)** The inner color in orange shows the genes significantly associated with the expression of psychomotor agitation/retardation. The surrounding circle shows if the gene is differentially expressed in male (blue), female (pink) or both (dark violet). Hubs and nodes are defined by the size of the circles. **(b,f)** Correlation between module membership (KME) and gene significance values of the association between each gene and relevant symptomatic feature, for genes co-expressed in **(b)** *Darkorange* and **(f)** *Saddlebrown,* in males (blue) and females (pink) with the 95% confidence interval bands. **(c,g)** Pie chart showing the number of genes in **(c)** *Darkorange* associated with change in appetite/weight in green and **(g)** *Saddlebrown* associated with psychomotor agitation/retardation in orange. **(d,h)** GO enrichment for **(d)** *Darkorange* and **(h)** *Saddlebrown* modules for the first five most significantly enriched GO terms. The black vertical line indicates the significance threshold (p_adj_ <0,05).

Finally, we highlighted a small fraction of gene networks associated with the same symptomatic features in both sexes (**Fig. 6a**; **Supp. Table 10**). In total, we identified 22 gene modules associated with the same symptoms in males and females. However, 17 (77%) showed opposite associations in males and females and divergent genes responsible for these association. For instance, the expression of the *Orange* module in the OFC (**Fig. 6b**), enriched in genes involved in synaptic transmission (p_adj_<5.0^E-9^), is positively correlated with the expression of difficulty in concentration/indecision in males with MDD (r=0.484, p_adj_<0.05), but negatively correlated with the same symptom in female MDD (r=-0.622, p_adj_<0.05) (**Fig. 6c**). Furthermore, out of the genes significantly associated with the expression of this symptom in both sexes (51 in male MDD and 46 in female MDD), only six were common in males and females (**Fig. 6d**). Noticeably, only one hub gene (*Ppp3r1*) was commonly associated with difficulty in concentration/indecision in males and females with MDD, with all other hub genes being either uniquely associated with this clinical feature in males (*Gabrb3*, *Prkce*) or females (*Synj1*, *Atp9a*, *Snap91*, *Rab6b*) with MDD (**Fig. 6b**). Similarly, the expression of *Lightcyan1* in the NAc (**Fig. 6e**), enriched with genes relevant to function of the mitochondria (p_adj_<5.^0E-11^), was negatively correlated with the expression of insomnia/hypersomnia in male MDD (r=-0.409, p_adj_=0.078), but positively correlated with the same symptom in females with MDD (r=0.875, p_adj_<0.05; **Fig. 6f**). *Lightcyan1* in NAc is also enriched for genes upregulated in male (p_adj_<0.01) but not female MDD. Six (*Gpx4*, *Psmb5*, *Psmb6*, *Prdx5*, *Asna1*, *Eif4h*) out all seven hub genes were associated with the expression of insomnia/hypersomnia only in females, while *Urod* was common to both sexes (**Fig. 6e, g**). Together, these findings suggest that while these networks may act as ensembles of genes underlying the expression of clinical features of MDD in both sexes, their specific associations in either males or females may be driven largely by different gene members.

**Figure 6.**
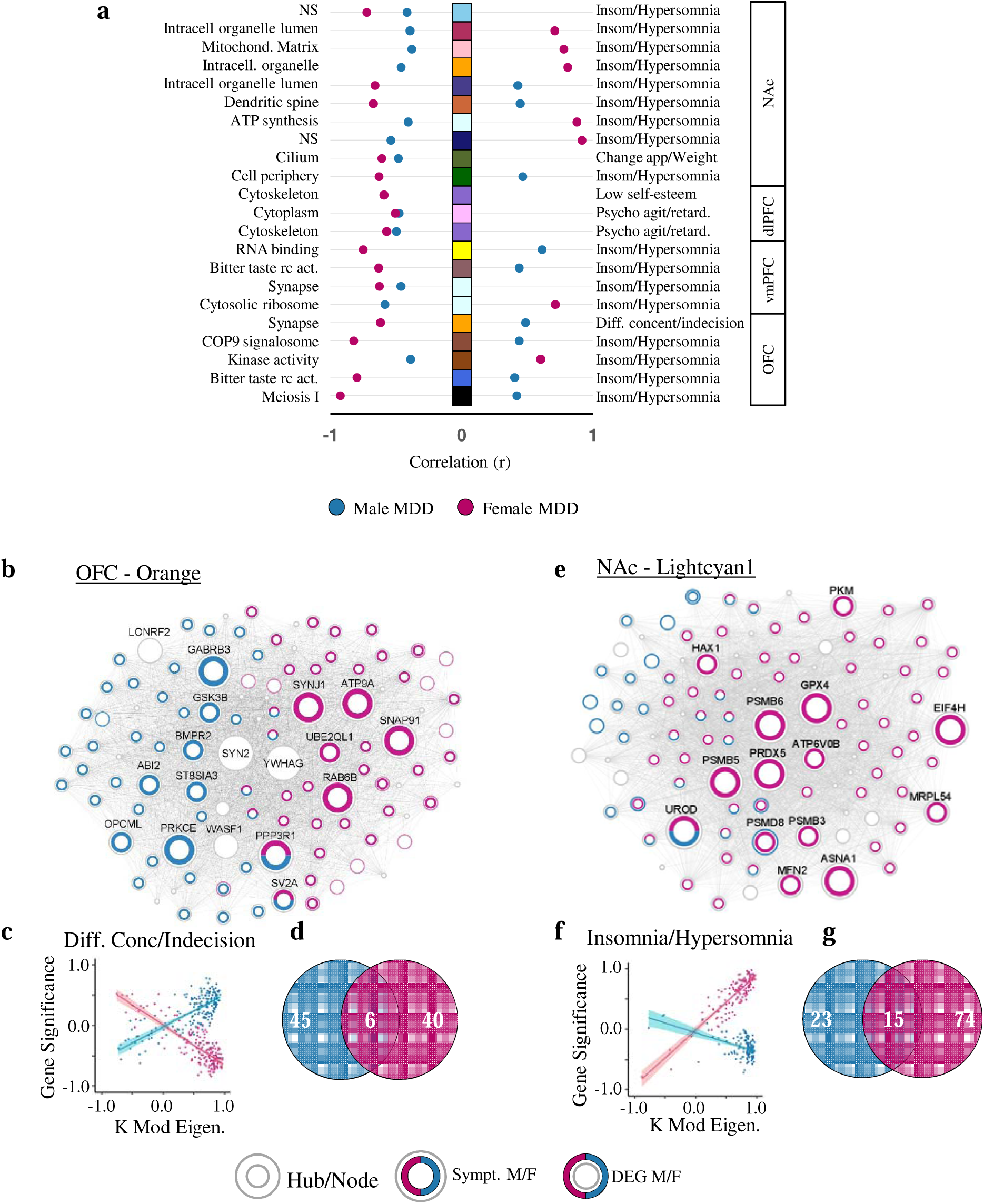
Common symptoms in males and females are driven in part by the activity of different genes. **(a)** List of gene networks associated with the same symptomatic features in both sexes, with their respective GO terms on the left, and symptoms across brain regions on the right. The squares in the middle show modules with their assigned arbitrary names (colors). The dots show the correlation values between module membership (KME) and gene significance values, with blue for males and pink for females. **(b,e)** Hubs and nodes representation of **(b)** module *Orange* in the OFC, showing less than 4% of the genes in this module commonly associated with the expression of difficulty in concentration/indecision in males and females. **(e)** Module *Lightcyan1* in the NAc in which 10% of the genes are commonly associated with the expression of insomnia/hypersomnia in males and females. **(c,f)** Correlations between module membership (KME) and gene significance values of the association between each gene and relevant symptomatic feature, for genes co-expressed in **(c)** module *Orange,* and **(f)** module *lightcyan1* in males (blue) and females (pink) with the 95% confidence interval bands. Opposite direction of this association in males and females reveals an opposite structural association in the two sexes. **(d,g)** Venn diagrams showing the number of genes in **(d)** module *Orange* associated with difficulty in concentration/indecision and **(g)** module *Lightcyan1* associated with insomnia/hypersomnia in males (blue), females (pink), or in common in both sexes.

## Discussion

For decades, the expression of MDD’s clinical manifestations have been related to variations in the activity of specific brain functions^57–61^, while the molecular mechanisms underlying these changes have never been fully explored. Here, we provide a thorough and unbiased description of transcriptional signatures across brain regions associated with the expression of MDD and, more specifically, with the expression of some of its core symptom domains in males and females. Our results suggest that the expression of specific symptoms results from the activity of different gene networks across brain regions. Our findings point toward specific brain structures as being more relevant for the expression of the different symptom domains in males and females. Moreover, although the transcriptional organization of gene networks may be preserved in both sexes, their association with the expression of its symptoms differs significantly in males versus females.

MDD in males and females is defined through the same clinical criteria and both sexes express the same symptoms although at somewhat different levels^62,63^. Accordingly, one may expect similar molecular mechanisms underlying the expression of these symptoms in both sexes. However, our results suggest otherwise, even though the transcriptional organization of gene networks are strongly conserved in both sexes. Indeed, we only identified a small proportion of modules associated with the expression of the same symptoms in males and females and, for most of them, we saw opposite correlations between gene networks and symptom in males versus females, consistent with a prior report^22^. For instance, we identified the *Orange* module in the OFC and *Lightcyan1* module in the NAc associated with the expression of difficulty in concentration/indecision and insomnia/hypersomnia, respectively, in both males and females. These two modules are enriched in genes involved in synaptic transmission and function of the mitochondria, two fundamental processes in males and females that have been implicated previously in MDD^26–29^. However, while this may support the idea of common functional and behavioral implications for these two gene networks in male and female symptomatology, it also suggests that these genes may be acting differently in both sexes. This is further supported by our findings showing that the genes significantly associated with the expression of these symptoms in males and females with MDD were drastically different. In the OFC, hub genes in the *Orange* module encoding the glycogene synthase kinase 3 (*Gsk3b*) and the GABA receptor subunit ß3 (*Gabrb3*) were specifically associated with the expression of difficulty in concentration/indecision in male MDD, while hub genes encoding synpatojanin 1 (*Synj1*) and the clathrin coat assembly protein AP180 (*Snap91*) drove the same associations in women. These genes may modulate neurotransmission differently in males versus females in pathological contexts. Overall, while the precise mechanisms underlying these effects remain to be elucidated, our findings further strengthen the hypothesis^28^ that common functional pathways are affected in both males and females with MDD, although through the action of different genes, and expand this concept to specific symptomatology of MDD.

Changes in network structures found in specific brain regions may interfere with regional activity and consequently with the functional connectivity of brain networks controlling specific behavioral domains relevant to the expression of each symptom in males and females^7,8^. Here, although we did not empirically confirm the functional roles of our predicted gene networks, converging evidence in mouse models of chronic stress suggest that this may be the case^24,28,53,54^. For instance, the *Ivory* gene network in the OFC, associated with the expression of three main symptoms of MDD in males, is enriched for genes relevant to GABAergic neurotransmission. Alterations of the GABAergic system are a hallmark of MDD and have been associated with disruption of the homeostatic inhibitory control over excitatory tone that is required for the top-down processing of cognitive and emotional information in cortical regions^55,56^. As part of a larger brain network, the OFC is hypothesized to be part of the attention and cognitive control circuitry, with alterations of this circuit causing indecision and decreased concentration and attention^4,15–17^. Similarly in females, we identified gene networks associated with the expression of change in appetite/weight (*Darkorange*) and low self-esteem (*Purple*) in the aINS, psychomotor agitation/retardation (*Saddlebrown*) in the dlPFC and difficulty in concentration/indecision (*Palevioletred3*) in the vmPFC that could also associate with changes in the activity of each of these brain regions. It is likely that changes in these sex-specific gene networks underly symptom expression in males or females by interfering with the activity of brain networks, as was recently shown for a cortical-subcortical circuit during adolescent development^64^. Thus, we hypothesize that the reorganization of precise transcriptional structures across brain regions in males and females may underly the expression of distinct clinical features of MDD in a sex-specific fashion.

Our approach in this study was to break down the complexity of MDD to its simplest expression, i.e., at the symptom level. However, various symptoms often co-occur. This is well exemplified through the clustering approaches we used at the gene level but could not be accounted for at the gene network level. Indeed, although we analyzed one of the largest available RNA-seq datasets for MDD in males and females, we did not reach sufficient power to evaluate the co-occurrence of specific symptoms in males and females at the network level. Similarly, the postmortem nature of our study does not allow us to evaluate the intensity and recurrence of each symptom in both sexes. Nevertheless, each of our predictions resulted from the sum of converging evidence cumulated from several levels of analysis. Furthermore, our analyses confirmed the reproducibility of our transcriptional observations, further supporting the validity of our findings.

To conclude, the heterogeneity of MDD has long been a brake toward an understanding of its molecular etiologies. Progress in computational biology combined with improvement in the collection of clinical information allowed us to transition from global transcriptional screenings^20–23^ to state versus trait transcriptional assessment^29^ and sex-specific transcriptional structures^26–28^ and ultimately toward the dissection of transcriptional signatures associated with symptomatic profiles in males and females. Findings from this study suggest such associations exist and strongly support the implementation of systems biology approaches to larger longitudinal cohorts with evolving pathological states. Furthermore, converging transcriptional signatures have been identified across several psychiatric conditions^27,65^. Dissecting the transcriptional signatures underlying the expression of clinical manifestations either common or unique to these conditions will further improve our capacity to diagnose each condition with greater precision and ultimately transition from the treatment of a syndrome toward the treatment of specific symptoms or disease states.

## Supporting information

Supplementary Table 3_PCA results by brain region in the 95 percent most variable genes

Supplementary Table 5_list of DEGs in men

Supplementary Table 6_list of DEGs in women

Supplementary Table 7_Gene Ontology Results of DEGs across brain regions in men

Supplementary Table 8_Gene Ontology Results of DEGs across brain regions in women

Supplemental Table 10_List of modules expressing common symptoms in men and women across six brain region

## Acknowledgments

BL holds a Sentinelle Nord Research Chair, is supported by the Canadian Institutes of Health Research (Grant No. PJT-451728 and PJT-451858), and the Natural Science and Engineering Research Council of Canada (Grant No. RGPIN-2019-06496) and receives Fonds de Recherche en Santé du Québec (FRQS) Junior-2 salary support. This work was also supported by the U.S. National Institute of Mental Health (Grant No. R01MH129306 to EJN).

## Author contributions

S.M. and B.L. conceived the project, designed the experiments and analyses, and wrote the manuscript. B.L. generated the data. T.H.C. contributed to the design and to the analyses. S.M.,

A.M.P. and A.M.R. analyzed all the data. All authors contributed to the preparation of the manuscript.

## Supplemental Figure legends

**Supplemental Figure 1.**
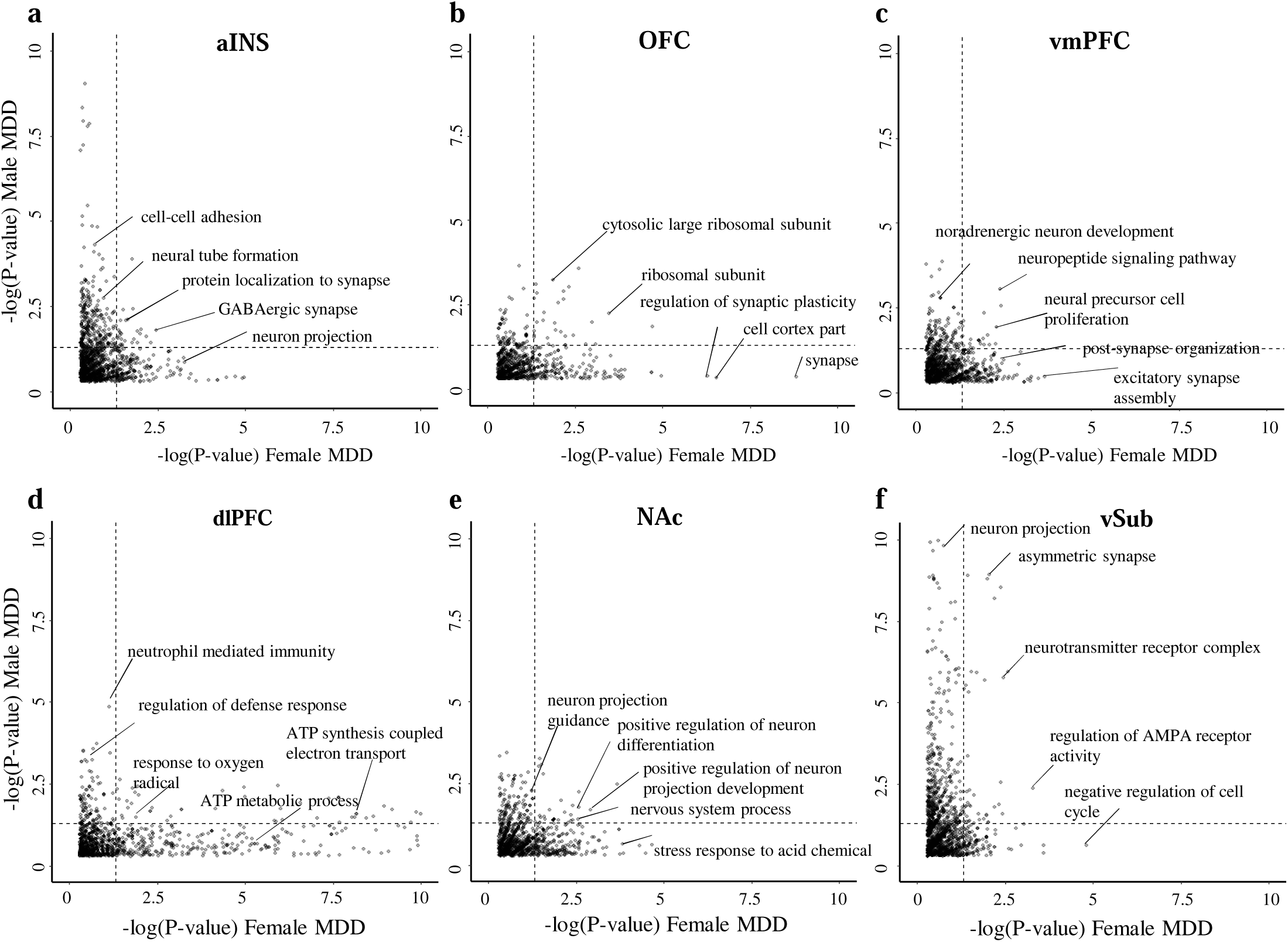
Common functional terms enriched in DEGs in males and females. (**a-f)** Scatter plots showing the ontological terms (commonly/differently) enriched for DEGs in males and females with MDD across brain regions.

**Supplemental Figure 2.**
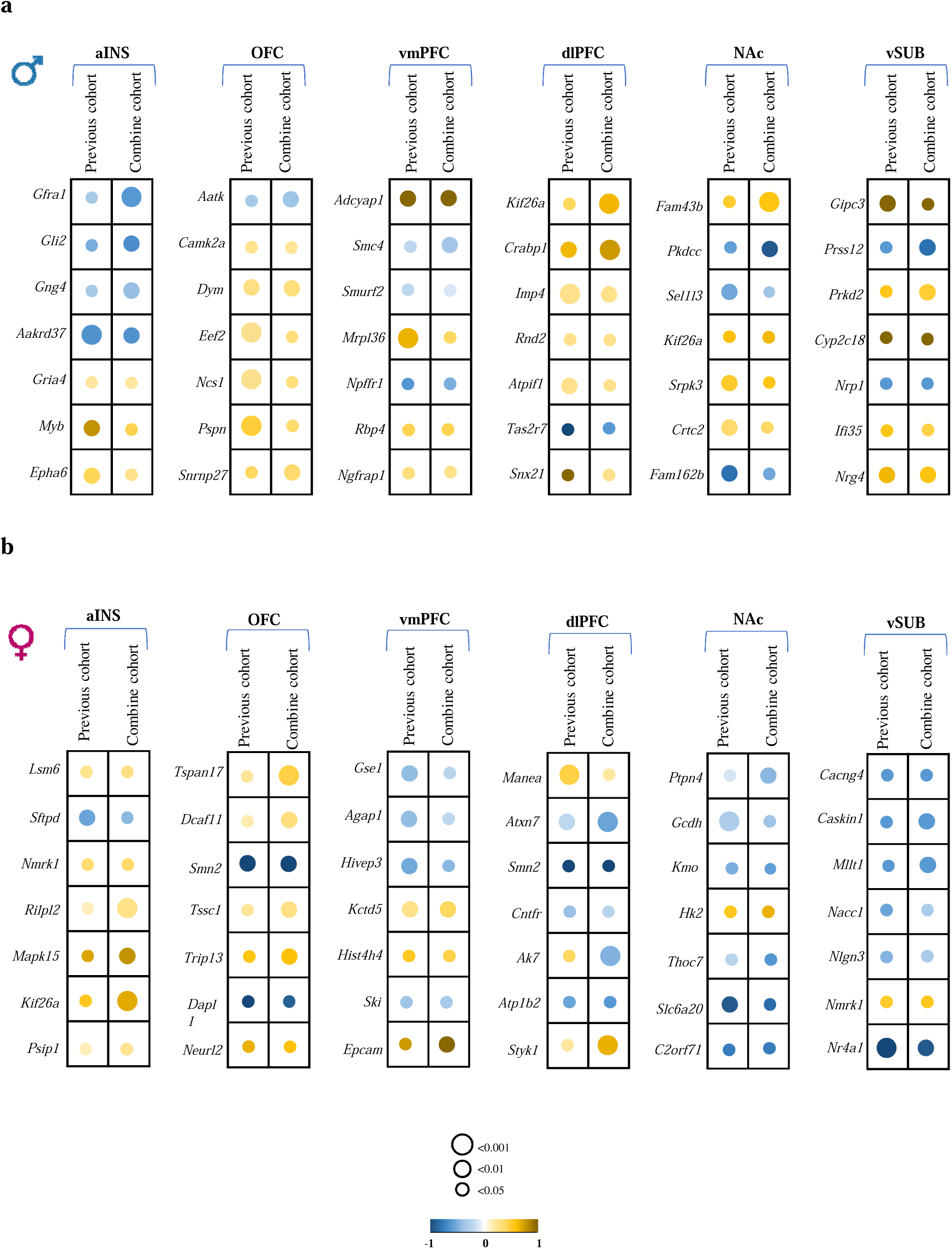
Comparison of DEGs from distinct postmortem cohorts shows consistent transcriptional changes in males and females across brain regions. Representative list of genes differentially expressed in this study and our previously published study^27^ in males (**a)** and females **(b)** across six brain regions. Colors represent log fold change values, with blue for genes downregulated and yellow for genes upregulated in MDD versus control conditions.

**Supplemental Figure 3.**
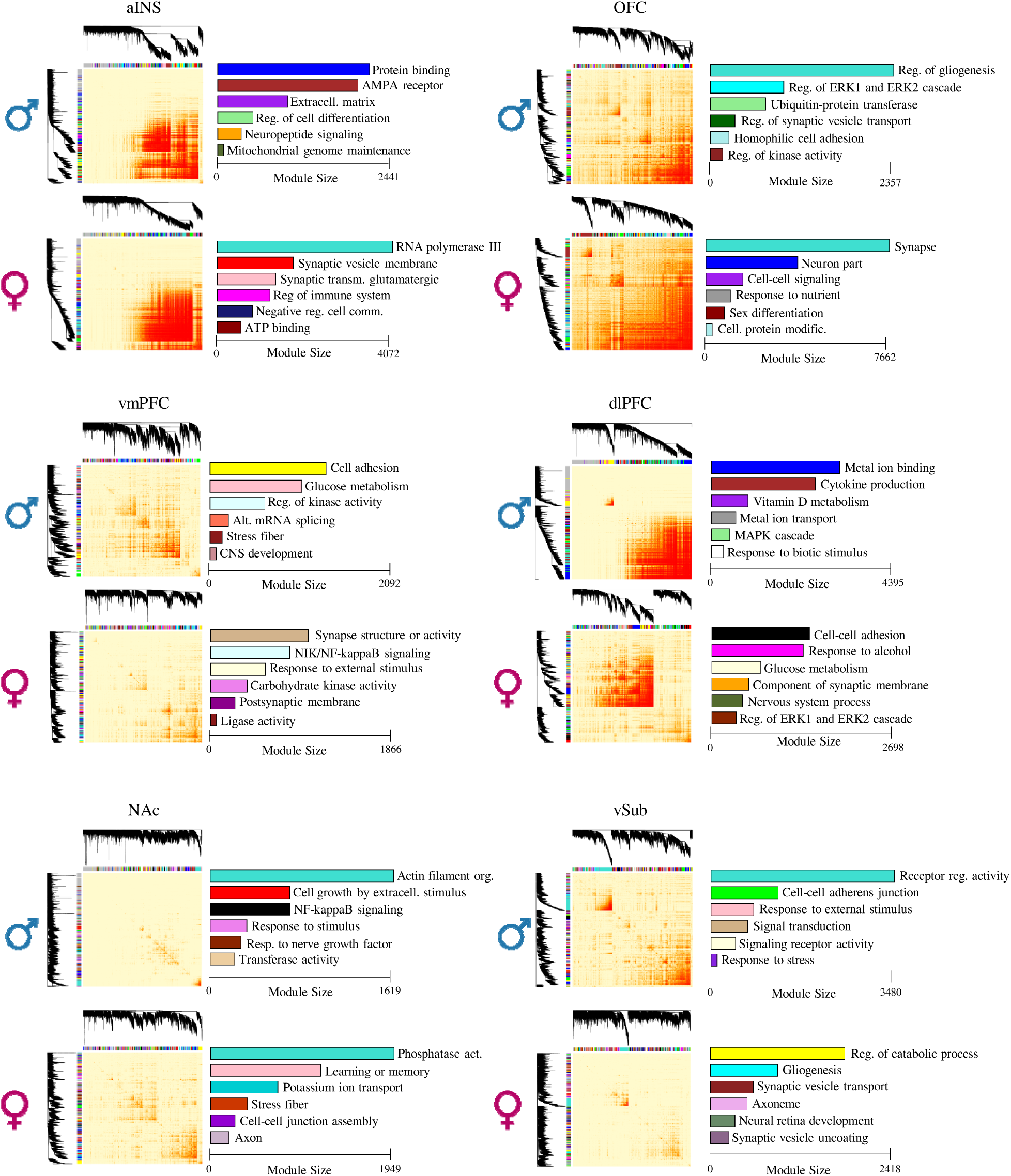
Topological overlap matrix (TOM) plots for MDD modules in males and females across six brain regions. The color bars at the right side show module size with their enriched ontological terms.

**Supplemental Figure 4.**
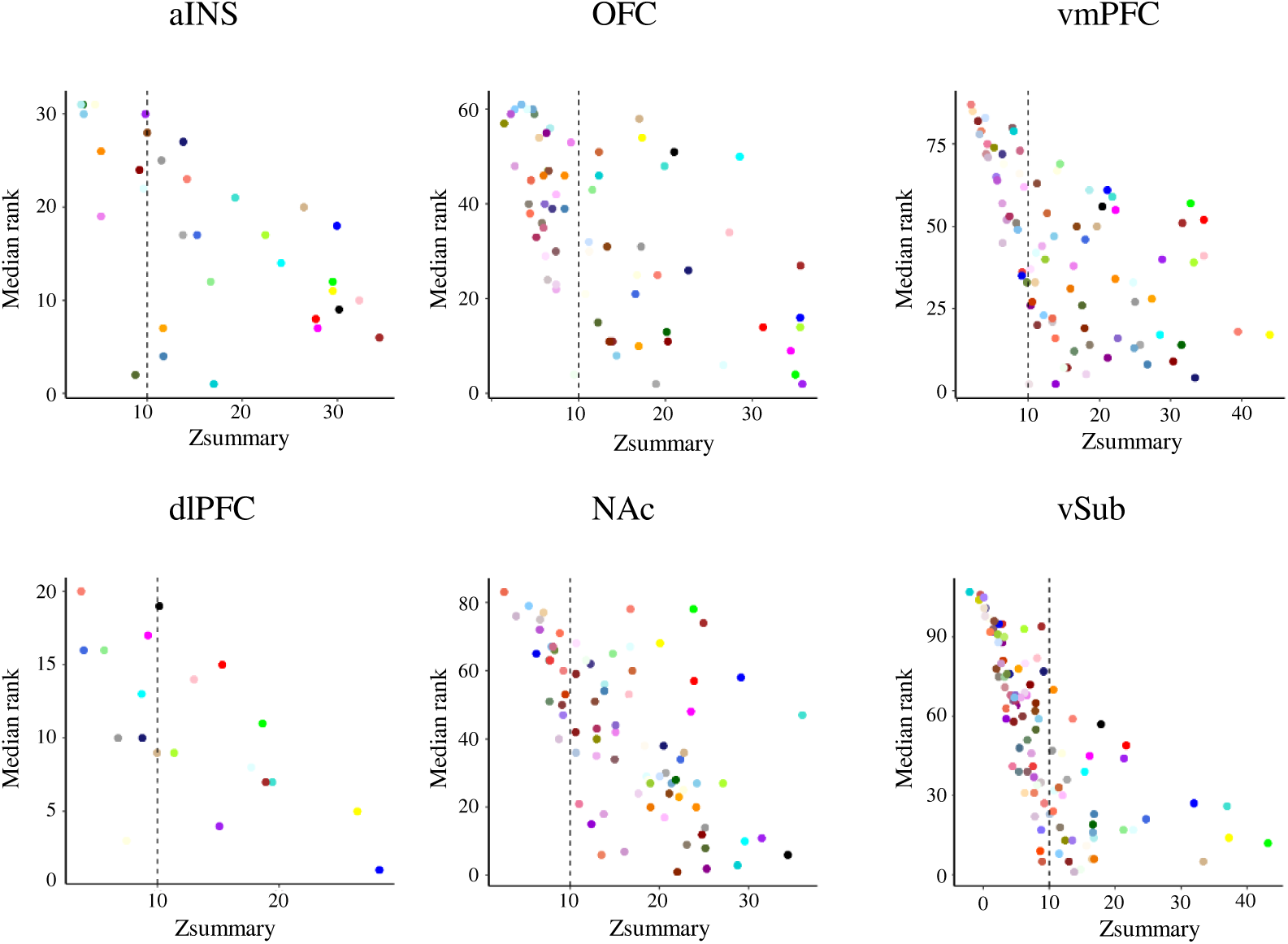
Preservation of the transcriptional organization of gene networks between male MDD and female MDD. Modules with Z_summary_ scores higher than 10 are considered as male MDD modules preserved in female MDD.

**Supplemental Figure 5.**
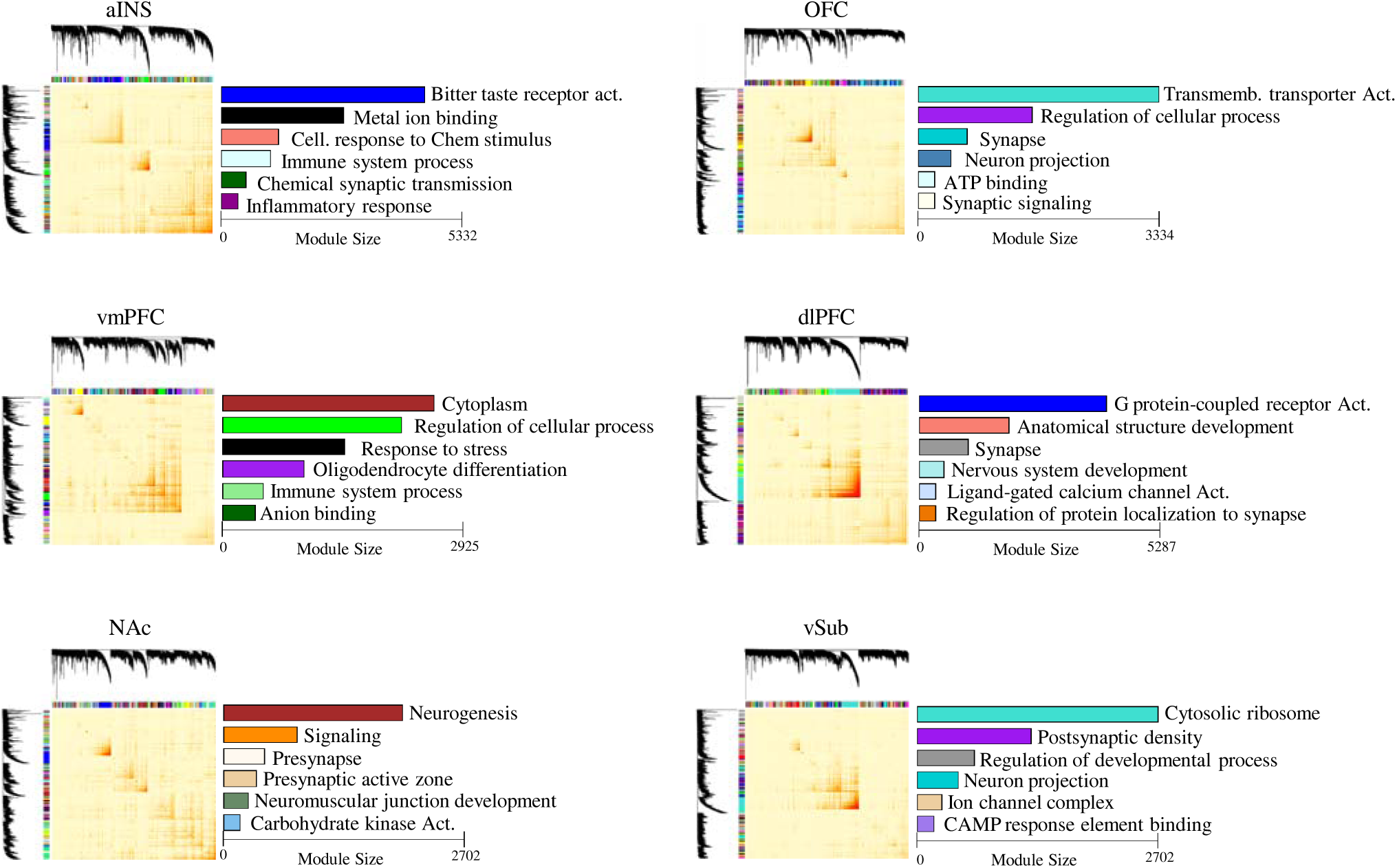
Topological overlap matrix (TOM) plots for modules combining males and females across six brain regions. The color bars at the right side show module size with their enriched ontological terms.

**Supplementary Figure 6.**
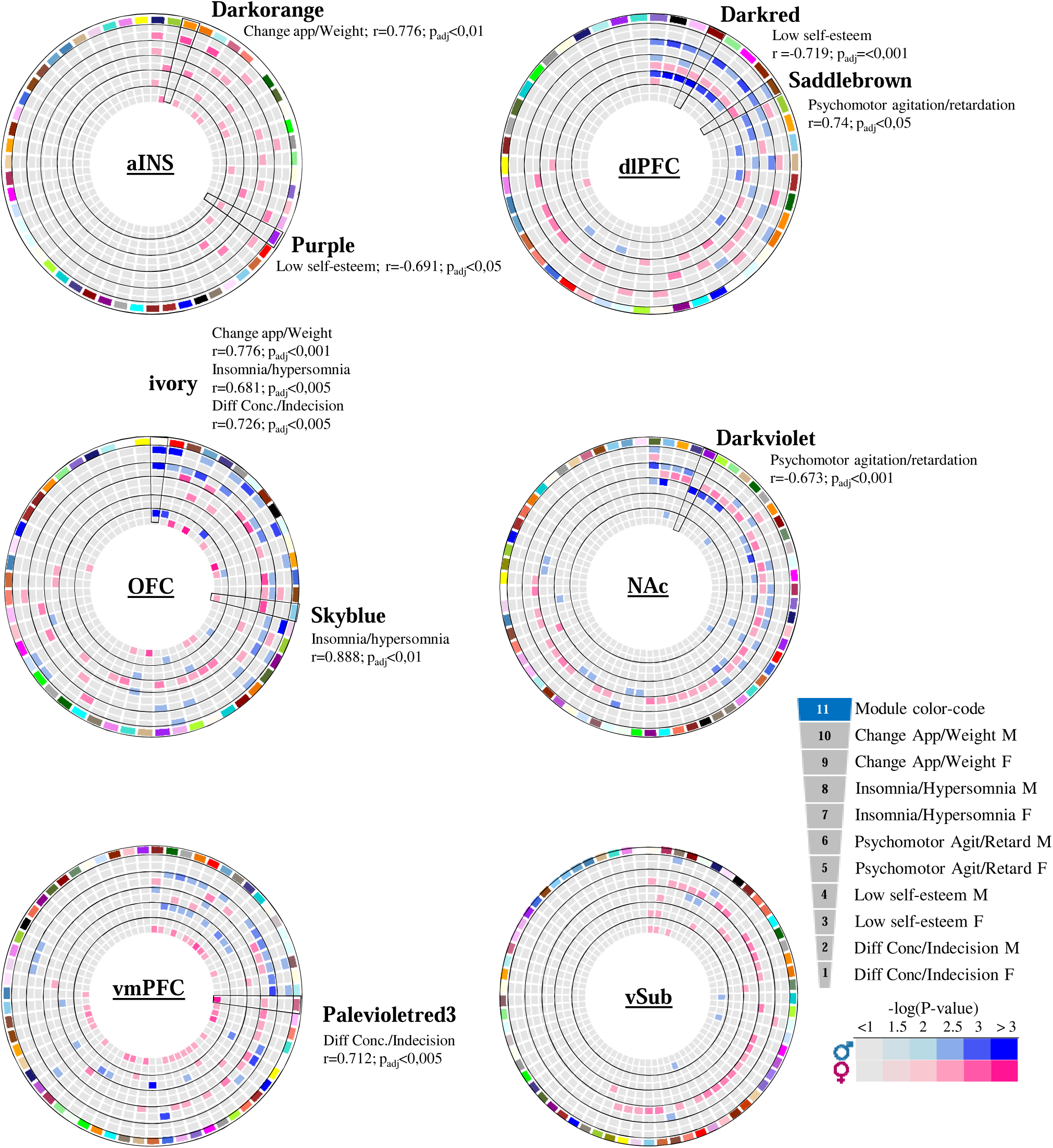
Circos plots displaying the degree of associations between modules and symptomatic features of MDD across six brain regions in males (blue) and females (pink). The key at the bottom of the circos plots represents the strength of each association. The outermost circle is an arbitrary color randomly assigned as the module name. The order of the symptom associations is, from top to bottom, change in appetite/weight, insomnia/hypersomnia, psychomotor agitation/retardation, low self-esteem and difficulty in concentration/indecision in male and females, respectively.

**Supplementary Figure 7.**
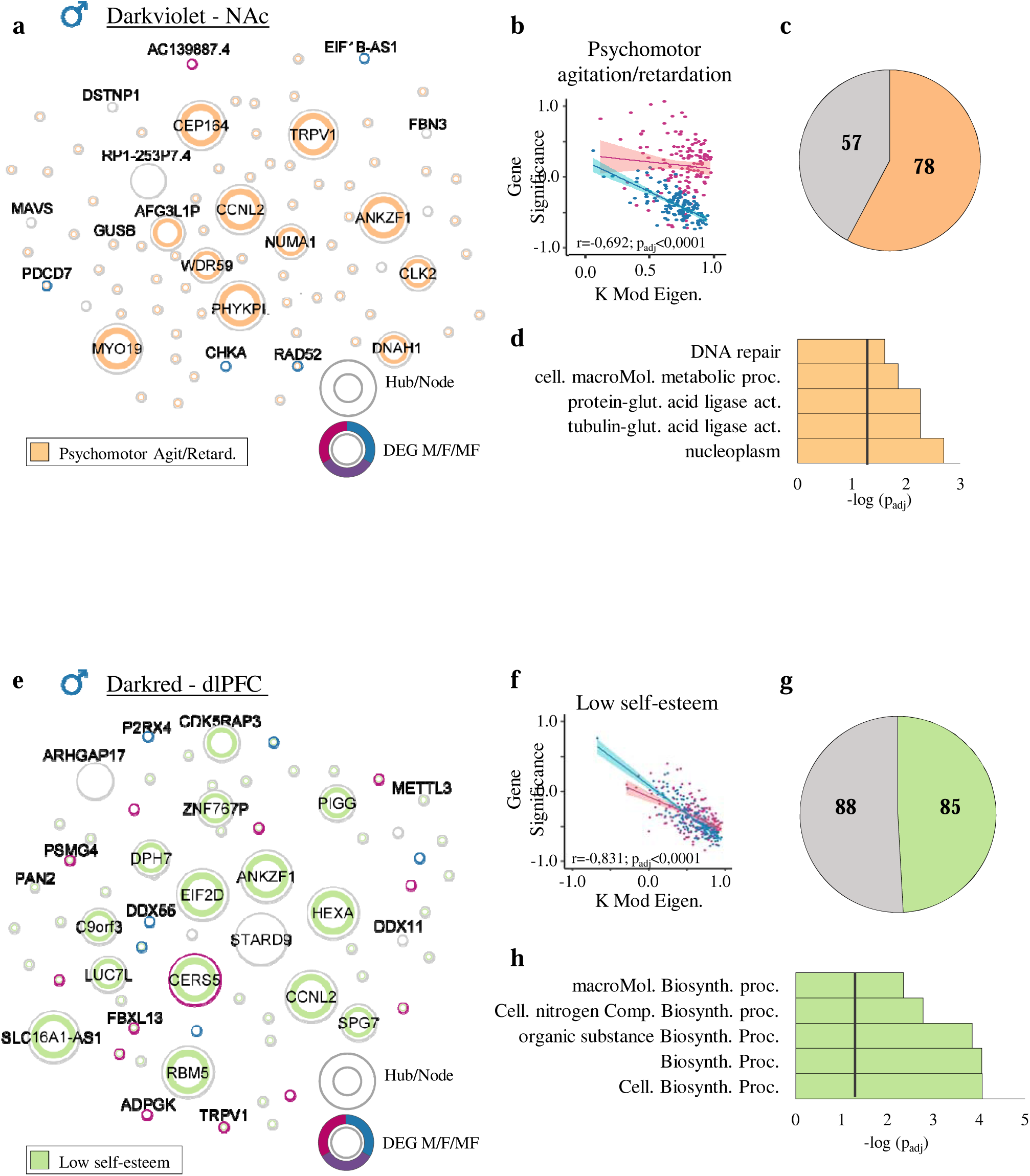
Representative modules (*Darkviolet* and *Darkred*) associated with the expression of psychomotor agitation/retardation and low self-esteem in male MDD identified in the NAc and dlPFC, respectively. **(a,e)** Hubs and nodes representation of **(a)** module *Ivory* and **(e)** module *Darkred* and their associations with respective clinical symptoms. **(a)** The inner color in orange shows the genes significantly associated with the expression of the symptom: psychomotor agitation/retardation. **(e)** The inner color in bright green shows the genes significantly associated with the expression of low self-esteem. The surrounding circle shows if the gene is differentially expressed in males (blue), females (pink) or both (violet). Hubs and nodes are defined by the size of the circles. **(b,f)** Correlation between module membership (KME) and gene significance values of the association between each gene and relevant symptomatic feature for genes co-expressed in **(b)** module *Darkvioloet* and **(f)** module *Darkred,* in males (blue) and females (pink) with the 95% confidence interval bands. **(c,g)** Pie charts showing the number of genes in **(c)** module *Darkviolet* associated with psychomotor agitation/retardation in orange and **(g)** module *Darkred* associated with low self-esteem in bright green. **(d,h)** GO enrichment for **(d)** module *Darkviolet* and **(h)** module *Darkred* for the first five most significantly enriched GO terms. The black vertical line indicates the significance threshold (p_adj_<0,05).

**Supplementary Fig. 8.**
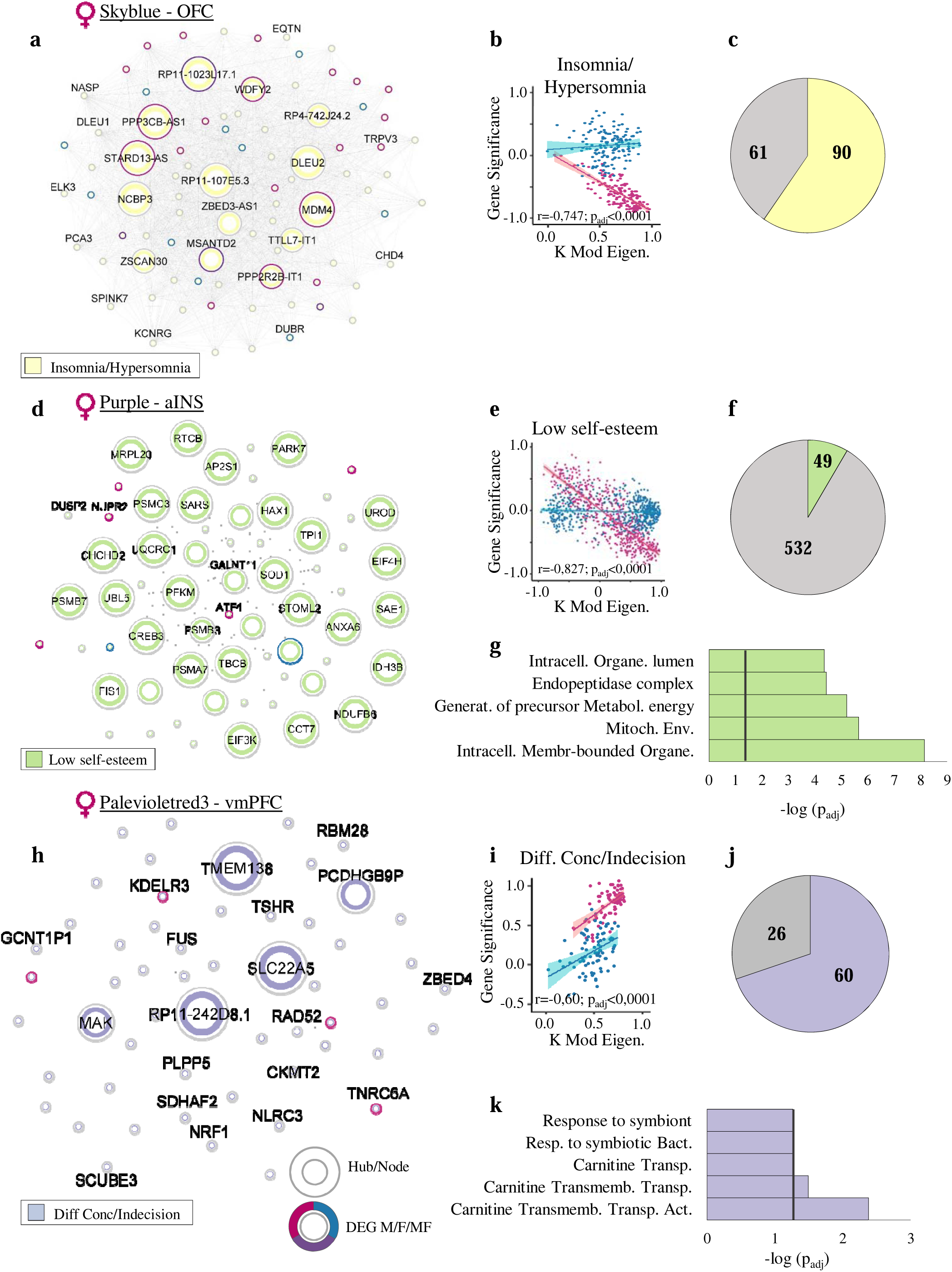
Representative modules (*Skyblue, Purple,* and *Palevioletred3*) associated with the expression of insomnia/hypersomnia, low self-esteem and difficulty in concentration/indecision in females detected in the OFC, aINS and vmPFC, respectively. **(a,d,h)** Hubs and nodes representation of **(a)** module *Skyblue,* **(e)** module *Purple*, (**h**) module *Palevioletred3* and their associations with respective clinical symptoms of MDD. The inner color shows the genes significantly associated with the expression of insomnia/hypersomnia (**a**, yellow), low self-esteem (**e**, green) and difficulty in concentration/indecision (**h**, purple). The surrounding circle shows if the gene is differentially expressed in males (blue), females (pink) or both (violet). Hubs and nodes are defined by the size of the circles. **(b,e,i)** Correlation between module membership (KME) and gene significance values of the association between each gene and respective symptomatic feature for genes co-expressed in **(b)** module *Skyblue,* **(e)** module *Purple* and **(i)** module *Palevioletred3* in males (blue) and females (pink) with the 95% confidence interval bands. **(c,f,j)** Pie charts showing the number of genes in **(c)** module *Skyblue* associated with insomnia/hypersomnia showing in yellow color, **(f)** module *Purple* associated with low self-esteem showing in bright green color and **(j)** module *Palevioletred3* associated with difficulty in concentration/indecision showing in purple color. **(g,k)** GO enrichment for **(g)** module *Purple* and **(k)** module *Palevioletred3* for the first five most significantly enriched GO terms. The black vertical line indicates the significance threshold (p_adj_ <0,05).

## References

1. Collaborators, G.D.a.I. Global burden of 369 diseases and injuries in 204 countries and territories, 1990-2019: a systematic analysis for the Global Burden of Disease Study 2019. Lancet (London, England) 396, 1204–1222 (2020).

2. Burcusa, S.L. & Iacono, W.G. Risk for recurrence in depression. Clinical psychology review 27, 959–985 (2007).

3. American Psychiatric Association. Diagnostic and Statistical Manual of Mental disorders, (Washington, DC, 2013).

4. Bartova, L., et al. Reduced default mode network suppression during a working memory task in remitted major depression. Journal of psychiatric research 64, 9–18 (2015).

5. Jang, K.L., Livesley, W.J., Taylor, S., Stein, M.B. & Moon, E.C. Heritability of individual depressive symptoms. Journal of affective disorders 80, 125–133 (2004).

6. Guintivano, J., et al. Identification and replication of a combined epigenetic and genetic biomarker predicting suicide and suicidal behaviors. Am J Psychiatry 171, 1287–1296 (2014).

7. Goldstein-Piekarski, A.N., et al. Mapping Neural Circuit Biotypes to Symptoms and Behavioral Dimensions of Depression and Anxiety. Biological psychiatry 91, 561–571 (2022).

8. Williams, L.M. Precision psychiatry: a neural circuit taxonomy for depression and anxiety. The lancet. Psychiatry 3, 472–480 (2016).

9. Hamilton, J.P., Farmer, M., Fogelman, P. & Gotlib, I.H. Depressive Rumination, the Default-Mode Network, and the Dark Matter of Clinical Neuroscience. Biological psychiatry 78, 224–230 (2015).

10. Sheline, Y.I., Price, J.L., Yan, Z. & Mintun, M.A. Resting-state functional MRI in depression unmasks increased connectivity between networks via the dorsal nexus. Proceedings of the National Academy of Sciences of the United States of America 107, 11020–11025 (2010).

11. Matthews, S.C., Strigo, I.A., Simmons, A.N., Yang, T.T. & Paulus, M.P. Decreased functional coupling of the amygdala and supragenual cingulate is related to increased depression in unmedicated individuals with current major depressive disorder. J Affect Disord 111, 13–20 (2008).

12. Jaworska, N., Yang, X.R., Knott, V. & MacQueen, G. A review of fMRI studies during visual emotive processing in major depressive disorder. The world journal of biological psychiatry : the official journal of the World Federation of Societies of Biological Psychiatry 16, 448–471 (2015).

13. Mulders, P.C., van Eijndhoven, P.F., Schene, A.H., Beckmann, C.F. & Tendolkar, I. Resting-state functional connectivity in major depressive disorder: A review. Neuroscience and biobehavioral reviews 56, 330–344 (2015).

14. Peterson, A., Thome, J., Frewen, P. & Lanius, R.A. Resting-state neuroimaging studies: a new way of identifying differences and similarities among the anxiety disorders? Canadian journal of psychiatry. Revue canadienne de psychiatrie 59, 294–300 (2014).

15. Sylvester, C.M., et al. Functional network dysfunction in anxiety and anxiety disorders. Trends in neurosciences 35, 527–535 (2012).

16. Qiu, C., et al. Regional homogeneity changes in social anxiety disorder: a resting-state fMRI study. Psychiatry research 194, 47–53 (2011).

17. Korgaonkar, M.S., Grieve, S.M., Etkin, A., Koslow, S.H. & Williams, L.M. Using standardized fMRI protocols to identify patterns of prefrontal circuit dysregulation that are common and specific to cognitive and emotional tasks in major depressive disorder: first wave results from the iSPOT-D study. Neuropsychopharmacology : official publication of the American College of Neuropsychopharmacology 38, 863–871 (2013).

18. Treadway, M.T. & Zald, D.H. Reconsidering anhedonia in depression: lessons from translational neuroscience. Neuroscience and biobehavioral reviews 35, 537–555 (2011).

19. Kim, M.J., Hamilton, J.P. & Gotlib, I.H. Reduced caudate gray matter volume in women with major depressive disorder. Psychiatry research 164, 114–122 (2008).

20. Sequeira, A., et al. Global brain gene expression analysis links glutamatergic and GABAergic alterations to suicide and major depression. PLoS One 4, e6585 (2009).

21. Sequeira, A., et al. Implication of SSAT by gene expression and genetic variation in suicide and major depression. Arch Gen Psychiatry 63, 35–48 (2006).

22. Seney, M.L., et al. Opposite Molecular Signatures of Depression in Men and Women. Biological psychiatry 84, 18–27 (2018).

23. Sequeira, A., et al. Patterns of gene expression in the limbic system of suicides with and without major depression. Mol Psychiatry 12, 640–655 (2007).

24. Issler, O., et al. Sex-Specific Role for the Long Non-coding RNA LINC00473 in Depression. Neuron 106, 912–926.e915 (2020).

25. Bagot, R.C., et al. Ketamine and Imipramine Reverse Transcriptional Signatures of Susceptibility and Induce Resilience-Specific Gene Expression Profiles. Biol Psychiatry 81, 285–295 (2017).

26. Scarpa, J.R., et al. Shared Transcriptional Signatures in Major Depressive Disorder and Mouse Chronic Stress Models. Biological psychiatry In press (2020).

27. Girgenti, M.J., et al. Transcriptomic organization of the human brain in post-traumatic stress disorder. Nature neuroscience 24, 24–33 (2021).

28. Labonte, B., et al. Sex-specific transcriptional signatures in human depression. Nat Med (2017).

29. Shukla, R., et al. Molecular characterization of depression trait and state. Mol Psychiatry (2021).

30. Zhang, B., et al. Integrated systems approach identifies genetic nodes and networks in late-onset Alzheimer’s disease. Cell 153, 707–720 (2013).

31. Parikshak, N.N., et al. Integrative functional genomic analyses implicate specific molecular pathways and circuits in autism. Cell 155, 1008–1021 (2013).

32. Gandal, M.J., et al. Transcriptome-wide isoform-level dysregulation in ASD, schizophrenia, and bipolar disorder. Science (New York, N.Y.) 362 (2018).

33. Nolte, J. The Human Brain: An Introduction to Its Functional Neuroanatomy., (Mosby-Year Book Inc., St-Louis, MO, 2002).

34. McGirr, A., et al. Risk factors for completed suicide in schizophrenia and other chronic psychotic disorders: a case-control study. Schizophr Res 84, 132–143 (2006).

35. Dumais, A., et al. Risk factors for suicide completion in major depression: a case-control study of impulsive and aggressive behaviors in men. Am J Psychiatry 162, 2116–2124 (2005).

36. Spitzer, R.L., Williams, J.B., Gibbon, M. & First, M.B. The Structured Clinical Interview for DSM-III-R (SCID). I: History, rationale, and description. Arch Gen Psychiatry 49, 624–629 (1992).

37. Ritchie, M.E., et al. limma powers differential expression analyses for RNA-sequencing and microarray studies. Nucleic acids research 43, e47 (2015).

38. Smyth, G.K. Limma: linear models for microarray data. in Bioinformatics and Computational Biology Solutions using R and Bioconductor, Vol. 1 (ed. R. Gentleman, V.C., S. Dudoit, R. Irizarry, W. Huber) 397–420 (Springer, New York, 2005).

39. Risso, D., Ngai, J., Speed, T.P. & Dudoit, S. Normalization of RNA-seq data using factor analysis of control genes or samples. Nature biotechnology 32, 896–902 (2014).

40. Cahill, K.M., Huo, Z., Tseng, G.C., Logan, R.W. & Seney, M.L. Improved identification of concordant and discordant gene expression signatures using an updated rank-rank hypergeometric overlap approach. Scientific reports 8, 9588 (2018).

41. Plaisier, S.B., Taschereau, R., Wong, J.A. & Graeber, T.G. Rank-rank hypergeometric overlap: identification of statistically significant overlap between gene-expression signatures. Nucleic acids research 38, e169 (2010).

42. Benjamini, Y. & Yekutieli, D. The control of the false discovery rate in multiple testing under dependency. Ann Stat 29, 1165–1188 (2001).

43. Kolberg, L., Raudvere, U., Kuzmin, I., Vilo, J. & Peterson, H. gprofiler2 -- an R package for gene list functional enrichment analysis and namespace conversion toolset g:Profiler. F1000Research 9 (2020).

44. Langfelder, P. & Horvath, S. WGCNA: an R package for weighted correlation network analysis. BMC bioinformatics 9, 559 (2008).

45. Zhang, B. & Horvath, S. General framework for weighted gene coexpression analysis. in Statistical Applications in Genetics and Molecular Biology 4 (2005).

46. Song, W.M. & Zhang, B. Multiscale Embedded Gene Co-expression Network Analysis. PLoS computational biology 11, e1004574 (2015).

47. Smoot, M.E., Ono, K., Ruscheinski, J., Wang, P.-L. & Ideker, T. Cytoscape 2.8: new features for data integration and network visualization. Bioinformatics 27, 431–432 (2011).

48. Shannon, P., et al. Cytoscape: a software environment for integrated models of biomolecular interaction networks. Genome research 13, 2498–2504 (2003).

49. Shen, L. GeneOverlap: An R package to test and visualize gene overlaps. R Packag. (2014).

50. Camargo, A., Azuaje, F., Wang, H. & Zheng, H. Permutation - based statistical tests for multiple hypotheses. Source code for biology and medicine 3, 15 (2008).

51. Langfelder, P., Luo, R., Oldham, M.C. & Horvath, S. Is my network module preserved and reproducible? PLoS computational biology 7, e1001057 (2011).

52. Benjamini, Y. & Hochberg, Y. Controlling the False Discovery Rate - a Practical and Powerful Approach to Multiple Testing. Journal of the Royal Statistical Society Series B-Methodological 57, 289–300 (1995).

53. Bagot, R.C., et al. Circuit-wide Transcriptional Profiling Reveals Brain Region-Specific Gene Networks Regulating Depression Susceptibility. Neuron 90, 969–983 (2016).

54. Lorsch, Z.S., et al. Stress resilience is promoted by a Zfp189-driven transcriptional network in prefrontal cortex. Nature neuroscience 22, 1413–1423 (2019).

55. Duman, R.S., Sanacora, G. & Krystal, J.H. Altered Connectivity in Depression: GABA and Glutamate Neurotransmitter Deficits and Reversal by Novel Treatments. Neuron 102, 75–90 (2019).

56. Fogaça, M.V. & Duman, R.S. Cortical GABAergic Dysfunction in Stress and Depression: New Insights for Therapeutic Interventions. Frontiers in cellular neuroscience 13, 87 (2019).

57. Macey, P.M., et al. Brain Structural Changes in Obstructive Sleep Apnea. Sleep 31, 967–977 (2008).

58. Morrell, M.J., et al. Changes in brain morphology associated with obstructive sleep apnea. Sleep Medicine 4, 451–454 (2003).

59. Haase, L., Cerf-Ducastel, B. & Murphy, C. Cortical activation in response to pure taste stimuli during the physiological states of hunger and satiety. NeuroImage 44, 1008–1021 (2009).

60. LaBar, K.S., et al. Hunger selectively modulates corticolimbic activation to food stimuli in humans. Behavioral Neuroscience 115, 493–500 (2001).

61. Rolls, E.T. & Grabenhorst, F. The orbitofrontal cortex and beyond: From affect to decision-making. Progress in Neurobiology 86, 216–244 (2008).

62. Breslau, J., et al. Sex differences in recent first-onset depression in an epidemiological sample of adolescents. Transl Psychiatry 7, e1139 (2017).

63. Kessler, R.C. Epidemiology of women and depression. J Affect Disord 74, 5–13 (2003).

64. Dorfschmidt, L., et al. Sexually divergent development of depression-related brain networks during healthy human adolescence. Science advances 8, eabm7825 (2022).

65. Hartl, C.L., et al. Coexpression network architecture reveals the brain-wide and multiregional basis of disease susceptibility. Nat Neurosci 24, 1313–1323 (2021).

