## Supplementary tables: 1,2,4,9 for "Transcriptional dissection of symptomatic profiles across the brain of men and women with depression"

Supplementary Table 1. Postmortem samples demographics information

|  | Women CTRL | Women MDD | Men CTRL | Men MDD |
| --- | --- | --- | --- | --- |
| Age | 65.5±11.26 | 47.12±14.69 | 48.47±14.24 | 46.08±12.75 |
| PMI | 34.53±31.08 | 36.38±29.57 | 22.97±20.64 | 39.62±22.55 |
| pH | 6.38±0.41 | 6.64±0.3 | 6.49±0.33 | 6.65±0.3 |
| RIN aINS | 7.26±1.66 | 6.82±1.14 | 7.44±1.22 | 6.9±1.66 |
| RIN OFC | 7.46±1.52 | 7.26±0.84 | 7.5±1.09 | 7.18±1.45 |
| RIN vmPFC | 7.07±1.22 | 6.56±1.3 | 7.39±1.21 | 6.55±1.27 |
| RIN dlPFC | 7.31±1.14 | 7.1±0.84 | 7.59±1.21 | 7.25±1.19 |
| RIN NAc | 7.77±1.42 | 7.69±0.8 | 7.49±1.5 | 7.88±1.33 |
| RIN vSub | 6.27±1.45 | 6.88±1.04 | 6.88±1.79 | 7.33±1.51 |
| Childhood abuse  (NA/No/Yes) | 4/15/3 | 7/5/13 | 2/9/6 | 6/9/10 |
| Alcohol abuse  (NA/No/Yes) | 4/17/1 | 9/13/3 | 2/14/1 | 2/14/9 |
| Drug abuse  (NA/No/Yes) | 3/16/3 | 9/11/5 | 2/13/2 | 2/20/3 |
| Medication  (NA/No/Yes) | 4/15/3 | 4/4/17 | 1/15/1 | 5/9/11 |
| Smoking  (NA/No/Heavy/light/moderate) | 0/11/7/0/4 | 3/7/11/1/3 | 2/5/9/0/1 | 5/5/13/2 |
| Cohort  (Montreal/Texas) | 10/12 | 19/6 | 9/8 | 21/4 |

Supplementary Table 2. Frequency of Clinical symptoms in men and women with MDD across brain region

| Sex | Region | Response | Change in appetite/weight | Insomnia/  Hypersomnia | Psychomotor agitation/retardation | Low self-esteem | Difficulty in concentration/  indecision |
| --- | --- | --- | --- | --- | --- | --- | --- |
| Male MDD | aINS | No | 7 | 5 | 8 | 7 | 9 |
|  |  | Yes | 11 | 14 | 9 | 11 | 8 |
|  | OFC | No | 7 | 5 | 8 | 7 | 9 |
|  |  | Yes | 11 | 14 | 9 | 11 | 8 |
|  | vmPFC | No | 7 | 5 | 7 | 6 | 8 |
|  |  | Yes | 9 | 12 | 8 | 10 | 7 |
|  | dlPFC | No | 7 | 5 | 8 | 7 | 9 |
|  |  | Yes | 11 | 14 | 9 | 11 | 8 |
|  | NAc | No | 7 | 5 | 8 | 7 | 9 |
|  |  | Yes | 11 | 14 | 9 | 11 | 8 |
|  | vSub | No | 4 | 3 | 6 | 5 | 7 |
|  |  | Yes | 9 | 11 | 7 | 8 | 5 |
| Female MDD | aINS | No | 3 | 3 | 3 | 3 | 4 |
|  |  | Yes | 9 | 7 | 7 | 6 | 7 |
|  | OFC | No | 3 | 3 | 3 | 3 | 4 |
|  |  | Yes | 9 | 7 | 7 | 6 | 7 |
|  | vmPFC | No | 3 | 3 | 3 | 3 | 4 |
|  |  | Yes | 9 | 7 | 7 | 6 | 7 |
|  | dlPFC | No | 3 | 3 | 3 | 3 | 4 |
|  |  | Yes | 9 | 7 | 7 | 6 | 7 |
|  | NAc | No | 3 | 3 | 2 | 3 | 3 |
|  |  | Yes | 7 | 6 | 7 | 5 | 6 |
|  | vSub | No | 3 | 3 | 2 | 3 | 3 |
|  |  | Yes | 8 | 6 | 6 | 6 | 6 |

Supplementary Table 4. Cohort composition by brain region

| Region | Women CTRL | Women MDD | Men CTRL | Men MDD |
| --- | --- | --- | --- | --- |
| aINS | 21 | 23 | 16 | 25 |
| OFC | 21 | 23 | 16 | 24 |
| vmPFC | 19 | 25 | 17 | 22 |
| dlPFC | 21 | 24 | 17 | 24 |
| NAc | 19 | 22 | 15 | 24 |
| vSub | 12 | 19 | 12 | 18 |

Supplementary Table 9. Descriptive information of discovered modules in MDD group by sex and brain region

| Sex | Region | No. module | Min No. genes | Max No. Genes |
| --- | --- | --- | --- | --- |
| Male  MDD | aINS | 33 | 77 | 2441 |
|  | OFC | 67 | 50 | 2357 |
|  | vmPFC | 86 | 55 | 2092 |
|  | dlPFC | 20 | 122 | 4395 |
|  | NAc | 82 | 62 | 1619 |
|  | vSUB | 107 | 53 | 3480 |
| Female MDD | aINS | 25 | 87 | 4072 |
|  | OFC | 36 | 52 | 7662 |
|  | vmPFC | 78 | 51 | 1866 |
|  | dlPFC | 57 | 60 | 2698 |
|  | NAc | 105 | 51 | 1949 |
|  | vSub | 109 | 55 | 2418 |
